# Reelin doses normalizing behavioural alterations induced by repeated corticosterone avoid eliciting neuroinflammation-associated morphology in hippocampal microglia

**DOI:** 10.64898/2026.09.09.750432

**Authors:** Ciara Halvorson, Brady S. Reive, Sophie A. Thom, Callum M. Teevens, Josephine Han, Lisa E. Kalynchuk, Hector J. Caruncho

**Affiliations:** School of Medical Sciences, Faculty of Health, University of Victoria, Victoria, BC, Canada, V8P 5C2

## Abstract

Chronic stress elicits widespread, multisystem alterations, among them the increased production of inflammatory mediators by immune cells, that culminate in physiological and psychological disruptions. Microglia are resident macrophages of the CNS that show altered morphology alongside increases in pro-inflammatory cytokine release and phagocytosis in response to chronic stress. Chronic stress-induced behavioural and physiological alterations can be replicated in rats using daily injections of corticosterone (CORT) over a prolonged period. Reelin, a glycoprotein reduced in depression and animal models of chronic stress, rapidly reverses chronic CORT-induced symptoms following a single 3 μg injection. However, high levels of reelin may be pro-inflammatory, indicating the need to determine if a dose of reelin that resolves chronic stress-induced impairments exacerbates neuroinflammation as a possible adverse effect of reelin treatment. Using either a single 3 μg reelin injection following chronic stress or repeated (3) reelin injections throughout chronic stress in male Long Evans rats, we show that reelin does not promote microglia morphologies indicative of pro-inflammatory function within the context of chronic stress, and that both reelin administration schedules rapidly reverse chronic stress-induced despair-like behaviour. These results offer insight into the effect of peripheral reelin administration on processes underlying neuroinflammation and suggest that restoring reelin following chronic stress does not exacerbate microglia-driven neuroinflammation. This research together with past research from our lab supports the advancement of reelin or reelin peptides towards clinical evaluation as a rapid antidepressant.

## 1. Introduction

Chronic stress drives persistent glucocorticoid production and impacts proper functioning of the immune system. Physiological and behavioural alterations associated with long-term stress can be reliably reproduced in rats following daily injections of the glucocorticoid, corticosterone (CORT), over three weeks (Allen et al., 2022; Johnston et al., 2023). Reelin, a large (∼410 kDa) secreted extracellular matrix protein, reverses chronic CORT-induced alterations within one hour of administration, with effects persisting for up to one week (Scheil et al., 2024). Both individuals with depression and rodents exposed to chronic stress exhibit reduced levels of reelin in the plasma and hippocampus, while stressed rodents also show decreased reelin expression in the small intestine (Fatemi et al., 2000; Halvorson et al., 2025; Jin et al., 2022). Hippocampal reelin-immunoreactive (IR) cell density and other chronic CORT-associated alterations are normalized in rodents by traditional and non-traditional antidepressant interventions, including the administration of reelin (Brymer et al., 2018; Fenton et al., 2015; Johnston et al., 2023; Scheil et al., 2024). As chronic stress is a primary risk factor for major depressive disorder, and reelin exhibits a similar mechanism to the rapid antidepressant ketamine, reelin or reelin-related compounds present putative novel rapid-acting therapeutics for depressive disorders (Brymer et al., 2020; Johnston et al., 2023; Scheil et al., 2024; Slavich et al., 2009; Sullivan et al., 2000).

Although reelin injections exert antidepressant-like effects following chronic stress, high levels of reelin have been linked to inflammatory conditions. For example, blood reelin levels are elevated and correlated to pro-and anti-inflammatory markers in COVID-19 patients, and reelin depletion in rodent models of inflammatory conditions reduces pathological leukocyte infiltration to the central nervous system (CNS) during experimental autoimmune encephalitis (Calvier et al., 2020, 2023, 2024). Additionally, reduced reelin expression characterizes several neuropsychiatric and neurodegenerative disorders, including major depressive disorder, schizophrenia, Alzheimer’s disease, and autism, all of which also share chronic inflammation, as we’ve described previously (Calvier et al., 2023; Fatemi et al., 2000, 2005; Herring et al., 2012). These findings suggest that reelin homeostasis may be critical to immune and inflammatory homeostasis. However, the impact of peripheral reelin administration on inflammatory processes of the CNS in conditions of reduced reelin expression remains poorly understood.

Microglia, resident macrophages of the CNS, perform essential homeostatic functions, including synaptic remodeling, clearance of cellular debris and misfolded proteins, and support of neuronal health (Sierra et al., 2014). Microglial dysregulation has been increasingly linked to depression, where increased pro-inflammatory cytokine release, excessive engulfment of synaptic components, and diminished neurotrophic support converge to drive synaptic loss and impair both neurogenesis and neuronal plasticity (Kreisel et al., 2014). In response to chronic stress, microglia undergo morphological transformations towards ameboid or primed states that parallel these functional alterations. Ameboid microglia exhibit an enlarged soma, retracted processes, and reduced arborization, while primed microglia display an intermediate, sensitized phenotype between homeostatic (ramified) and ameboid microglia, showing an enlarged soma with reduced arbor complexity and thicker processes (Norden et al., 2015; Torres-Platas et al., 2014). Additionally, alterations in microglial density have been reported in stress-sensitive brain regions such as the hippocampus following chronic stress (Tong et al., 2017).

We recently showed multiple reelin injections modified the morphology of microglia in the CA3 region, however the changes did not reflect a clear shift towards a pro-or anti-inflammatory state - cell processes became more complex after reelin treatment in rats treated with chronic corticosterone and non-stressed controls (Reive et al., 2026). The present study investigates whether reelin treatments, which reversed corticosoterone-induced behavioural despair, altered microglial morphology within the polymorphic layer (PML) and subgranular zone (SGZ) of the hippocampus. In addition to evaluating microglia in additional regions, we evaluated the effects of a single injection of reelin against multiple injections of reelin and included both males and female rats. Lastly, we evaluated caspase-3 expression. Caspase-3 is a well-characterized executioner protease essential for apoptosis, but evidence suggests that caspase-3 activation is also involved in microglial regulation of neuroplasticity and neurogenesis, and microglia pro-inflammatory activation separately from its role in apoptosis (Alonso Bellido et al., 2023; Burguillos et al., 2011; Kavanagh et al., 2014).

## 2. Methods

### 2.1. Animal husbandry and chronic stress paradigm

Long-Evans rats (Charles River, Saint-Constant, QC, Canada) were singly housed in rectangular polypropylene cages in a colony room according to guidelines of the Canadian Council and Animal Care, and approval by the University of Victoria Committee on Animal Care. The room was maintained at 22+/- 1° C with a 12-hour light/dark cycle (lights on at 0700h and off at 1900h), and rats had unrestricted access to food and water. Rats were allowed to habituate for 7 days upon arrival and were handled briefly daily for 7 days prior to the commencement of the experiment. Thirty male and thirty female Long Evans rats were randomly assigned to one of five groups (n = 12; 6 males and 6 females per treatment group): Vehicle/Vehicle (V/V), Vehicle/Reelin (V/R), CORT/ Vehicle (C/V), CORT/Single Injection Reelin (C/R), or CORT/Repeated Reelin (C/RR). Rats received daily subcutaneous injections of either vehicle (V/V and V/R groups; 0.9% NaCl with 2% polysorbate-80), or CORT (C/V, C/R, and C/RR groups; 40 mg/ kg suspended in vehicle; Steraloids, Newport, RI) at a volume of 1ml/kg for 21 days. At three points over the experiment (days 1, 11, and 21), rats received an intravenous injection of either reelin or phosphate-buffered saline (PBS) through the lateral tail vein. V/V and C/V rats received 0.1M phosphate-buffered saline (PBS) on all three days. V/R and C/R rats received 0.1 M PBS on days 1 and 11, and recombinant reelin (3 μg, dissolved in 0.1 M PBS) on day 21. C/RR rats received recombinant reelin (3 μg, dissolved in 0.1 M PBS) on all three days. The recombinant reelin used consisted of the central reelin fragment (reelin repeats 3-6; 3820-MR-025-CF, R&D Systems), as this fragment elicits synaptic plasticity-related effects via canonical reelin receptor activation with comparable efficacy to the full-length reelin protein (Jossin, 2020; Li et al., 2023). Rats were weighed daily, weighing between 165-230 g (females) or 230-310 g (males) on the first day of injections (Fig. 2B-E).

### 2.2. Behavioural testing

#### 2.2.1. Forced swim test

We used a modified one-day forced swim test protocol as outlined in Allen et al., 2022. The containers used for the FST were rectangular Plexiglas containers of dimensions 25 cm wide x 25 cm long x 60 cm high, and were filled to a height of 30cm with water at 24 ± 1 °C. The testing took place between 2 and 4pm, approximately 6 hours post reelin injection. Rats were gently placed in the water and allowed to swim for 10 minutes. A video was recorded of each rat for future analysis of immobility. After swimming, rats were towel-dried and placed in their home cage under a heat lamp for approximately 20 minutes. Rats were tested one at a time. The tank was cleaned and the water was replaced following each rat. The videos were scored manually over the last 5 minutes of the test. Immobility was defined as no movement or only what is necessary to keep the head above water.

#### 2.2.2. Open field test

The open field test (OFT) was conducted one day after the FST. Each rat was placed facing the corner of an open-field arena (65 cm x 65 cm x 60 cm high) and allowed to explore the box for 10 minutes. Movement was monitored and automatically scored using EthoVision® tracking software (version 3.0, Wageningen, The Netherlands). Measurements were analyzed for time spent in the center zone (defined as 20 cm^2^ in the center of the box), frequency of center zone visits, average velocity, and total distance travelled. Between each rat, the box was sanitized using Virkon® (1%) disinfectant spray.

#### 2.2.3. Novel object location test

To assess spatial learning and memory, we used the novel object location test (NOLT). Rats were placed facing a corner of the same open field arena used in the OFT. In the training phase (4 mins), rats were allowed to explore two identical objects in diagonal corners of the box, leaving enough room so the rat could move behind the objects. Rats were removed from the box, and the box was sanitized with Virkon® (1%) disinfectant spray. The test phase took place 1 hour after the end of the training phase. In the test phase, one object was moved so that the objects were in adjacent corners of the box. Rats were placed in the same corner of the box as in the training phase and allowed to explore the two objects. After two minutes, rats were removed from the box and the box was sanitized using Virkon® (1%) disinfectant spray. A video of the NOLT was recorded using EthoVision® tracking software (version 3.0, Wageningen, The Netherlands). Videos were scored manually to obtain the amount of time spent exploring the moved object and the amount of time spent exploring the stationary object.

### 2.3. Tissue harvesting and processing

Rats were deeply anesthetized with isofluorane (5%) and perfused transcardially first with ice cold saline (0.9% NaCl) and then with 4% paraformaldehyde (PFA) once the blood was fully flushed from the body. Rats were decapitated and brains were collected and stored in 4% PFA for 12 hours. Brains were transferred to increasing concentrations of sucrose (10% *w/v* and 20% *w/v*) and stored in 30% sucrose (*w/v*) with 0.1% sodium azide.

Brains were embedded in optimal cutting temperature compound (OCT), and coronally cryosectioned at −20°C on a Leica Biosystems cryostat (CM1850 UV) at a thickness of 30 µm. Slices were stored at −20°C in cryoprotectant consisting of 30% *w/v* sucrose, 1% *w/v* PVP-40, and 30% ethylene glycol.

### 2.4. Immunohistochemistry

#### 2.4.1. Iba1

Every 12^th^ slice containing the dorsal hippocampus was selected for analysis, for a total of 3 slices (6 hippocampi) per rat. Free-floating sections were washed in 0.1M PBS, and quenched with 2% H_2_O_2_ diluted in 70% methanol for 10 mins and 0.1% NaBH_4_ diluted in PBS for 30 mins. After 3 washes in 0.1M PBS for 5 minutes each, sections were incubated in a blocking solution (10% FBS, 3% BSA (*w/v*), and 1% Triton X-100 in PBS) for 1 hour at room temperature (RT). Sections were incubated with an anti-Iba1 primary antibody (1:750 in blocking; Wako, 019-19741) overnight at 4°C. The next day, sections were washed in PBS (3 x 5 min), and incubated for 2 hours (RT) with a biotinylated goat anti-rabbit secondary antibody (1:300 in PBS; Vector Laboratories). Sections were washed in PBS (3 x 5 min). Sections were incubated in ABC (1:500 in PBS; Vector Laboratories) for 1 hour, washed in PBS (3 x 5 mins), and mounted onto Super-Frost Plus Microscope glass slides and allowed to dry. Slides were washed for 5 mins in sodium acetate (0.175M), and staining was revealed with 0.05% DAB (Sigma-Aldrich) and 0.015% H_2_O_2_ in 0.175M sodium acetate with 4.167% (*w/v*) nickel sulphate. Slides were serially dehydrated with ethanol and coverslipped with Permount mounting medium (Fisher Scientific SP15-500).

#### 2.4.2. Cleaved caspase-3

Every 12^th^ slice containing the dorsal hippocampus was selected for analysis, for a total of 3 slices (6 hippocampi) per rat. Free-floating sections were washed in 0.1M PBS, and incubated in a blocking solution containing 5% NGS, 1% BSA (*w/v*), and 0.3% Triton X-100 in PBS for 1 hour (RT). Sections were incubated overnight at 4°C with an anti-Iba1 primary antibody (1:200 in blocking; Abcam, ab283319) and an anti-cleaved-caspase-3 antibody (1:100; Cell Signaling, 9661S) diluted in blocking solution. The next day, sections were washed in PBS (3 x 5 min), and incubated for 1.5 hours (RT) in the secondary antibodies (1:200 in PBS; goat anti-rabbit ALEXA FLUOR 568, Invitrogen, A11011; goat anti-mouse ALEXA FLUOR 488, Invitrogen, A11001) at RT in the dark. Sections were washed in PBS (3 x 5 min). Sections were incubated with Hoestch (1:1000 in PBS) for 10 mins (RT), and washed in PBS (3 x 5 mins). Sections were mounted onto Super-Frost Plus Microscope glass slides, allowed to dry, and coverslipped using CitiFluor™ AF1 mounting medium (Electron Microscopy Sciences, 17970-100).

### 2.5. Microglia Density

Slide scanning images of the dorsal hippocampus were analyzed for microglia density in ImageJ. Microglia were counted in the PML and SGZ of the hippocampus, and regions were traced to obtain the density of microglia in both regions of the hippocampus. The PML was traced to obtain an area measurement (mm^2^), and the SGZ was traced with a line along the inside surface of the granular cell layer to obtain SGZ length (mm).

### 2.6. Analysis of microglia morphology

Various aspects of microglia morphology were considered using a variety of methodologies, as combining methods has been suggested elsewhere to ensure biologically accurate representations of microglia morphology (Green et al., 2022).

#### 2.6.1. Microglia process tracings

To evaluate whether chronic stress and/or treatment with reelin alters branching of hippocampal microglia, the processes of 6 cells per subject were manually traced at 63X magnification using Neurolucida software (version 2025.1.3, MBF Bioscience) to obtain a manually traced 3D skeleton representation of each microglia. Skeletons were flattened in Neurolucida Explorer (version 2025, MBF Bioscience) to obtain 2D representations of each microglia. Sholl analyses were conducted on microglia traces with Neurolucida Explorer to extract measures of intersections, nodes, endings and lengths at 10µm increments, with the center point at the center of the cell body, and totals for each measurement irrespective of distance.

#### 2.6.2. Microglia outline and soma tracings

Slides were imaged at 20x magnification using the slide scanning function on Neurolucida (version 2024.1.3, MBF Bioscience) to capture images of the full dorsal hippocampus for each brain slice. Slide-scanned images of the hippocampus were imported into ImageJ, where randomly selected microglia in the PML of the dentate gyrus (DG) were manually traced using the polygon tool with a screen-mirrored tablet and stylus for increased precision. 25 microglia outlines and 50 microglia soma were traced per rat in ImageJ. For SGZ tracings, slide-scanned images were imported into QuPath, where the wand tool was used to trace microglia. We transitioned from ImageJ to QuPath for image analysis, as tracing in QuPath was more time-efficient and comparison of tracings performed in ImageJ and QuPath demonstrated equivalent results in the PML (results not shown). Output measurements are defined in Figure 1 as follows: cell area, cell perimeter, circularity [(4π x area) / perimeter^2^], max feret (the longest distance between any two points along the tracing perimeter), aspect ratio (major axis/ minor axis, where major axis is the longest diameter of the best-fit ellipse, and the minor axis is the smallest diameter of the best-fit ellipse), roundness [(4 x area) / (π x major axis^2^)], and solidity (area / convex area; measures indentations in the cell boundary, or how “filled” the shape is).

**Figure 1:**
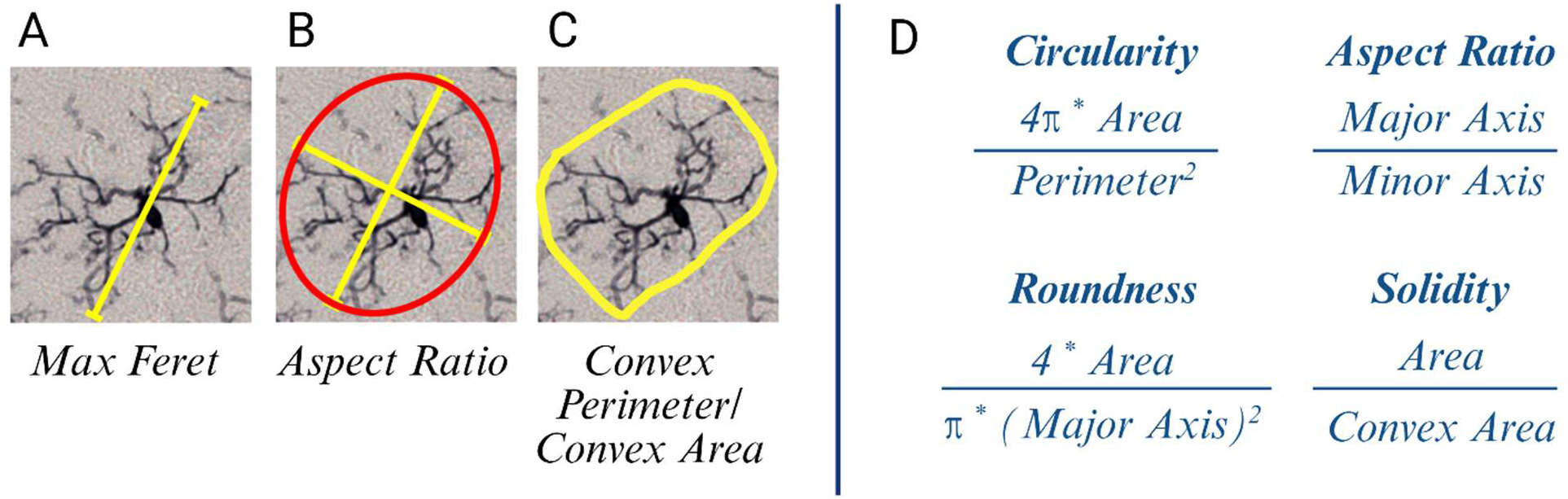
Visualization of morphological parameters. (A) Max feret refers to the shape’s maximum diameter, or the longest straight line across the microglia. (B) Aspect ratio measures a cell’s circularity and symmetry using the major and minor axes. (C) Convex perimeter and convex area are calculated from the cell’s convex hull (the smallest convex polygon that fully encloses the cell; analogous to an elastic band wrapping around the microglia’s outermost projections). Panel (D) shows the formulae for morphological parameters discussed in this paper.

### 2.7. Analysis of cleaved caspase-3 immunoreactivity in Iba1-immunoreactive microglia

Images of the dorsal hippocampus were taken at 20x magnification using the BFP, GFP, and dsRed channels (Neurolucida version 2025.1.3, MBF Bioscience). Images were loaded into ImageJ. At least 100 microglia per rat were counted and immunoreactivity for cleaved-caspase-3 (cleaved caspase-3) was determined in selected microglia in the PML and SGZ of the hippocampus. Data are expressed as the percentage of microglia (IR) for cleaved caspase-3 in each hippocampal layer.

### 2.8. Statistical analyses

Sample sizes were estimated on the basis of previous, similar studies. All data were collected by researchers blinded to the experimental condition. No data were excluded. For the analysis of daily weights, GraphPad Prism (version 10) was used to conduct a two-way repeated measures analysis of variance (ANOVA) (CORT x time) with Greenhouse–Geisser correction followed by Šídák’s multiple comparisons test for predefined groups. The NOLT parameters were analyzed with a three-way ANOVA (CORT x reelin x sex) followed by one-sample t-tests against chance (discrimination index (DI) of 0) in GraphPad. For the FST, OFT, microglia density, and cleaved caspase-3 measurements, we used three-way (ANOVAs) with treatment 1 (CORT or vehicle), treatment 2 (vehicle, reelin, or repeated reelin) and sex as independent factors. Šídák’s multiple comparisons test was conducted to perform pair-wise comparisons on predefined groups.

Outline tracings, process tracings, and soma area were analyzed using a nested linear mixed-effects model with stress, reelin, and sex as fixed effects and rat as a random intercept. Fixed effects were tested using Type III ANOVA. Significant effects were followed by Šídák-adjusted post hoc comparisons. Effect sizes for fixed effects were reported using R squared, and post hoc comparisons were quantified using Westfall’s d (Westfall et al., 2014).

GraphPad was used to create all graphs. All data is expressed as mean ± standard error of the mean (SEM), and significance was determined as p < 0.05. Full statistical results are available in Supplementary Table S1.

## 3. Results

### 3.1. CORT stunted weight gain in male rats but not in female rats

Daily injections of CORT hindered normal weight gain in male rats (Fig. 2B, C), with significant differences in vehicle/ CORT treatment group weights beginning by day 11 (p = 0.02) and continuing through the end of the experiment (p < 0.001). Female rats that received 21 days of CORT gained weight in a similar manner to vehicle rats (Fig. 2D, E). Male rats receiving vehicle gained significantly more weight than female rats receiving vehicle over the 21-day period (p < 0.001), and CORT altered weights more for male rats than females (Fig. 2F, G; p < 0.001).

**Figure 2.**
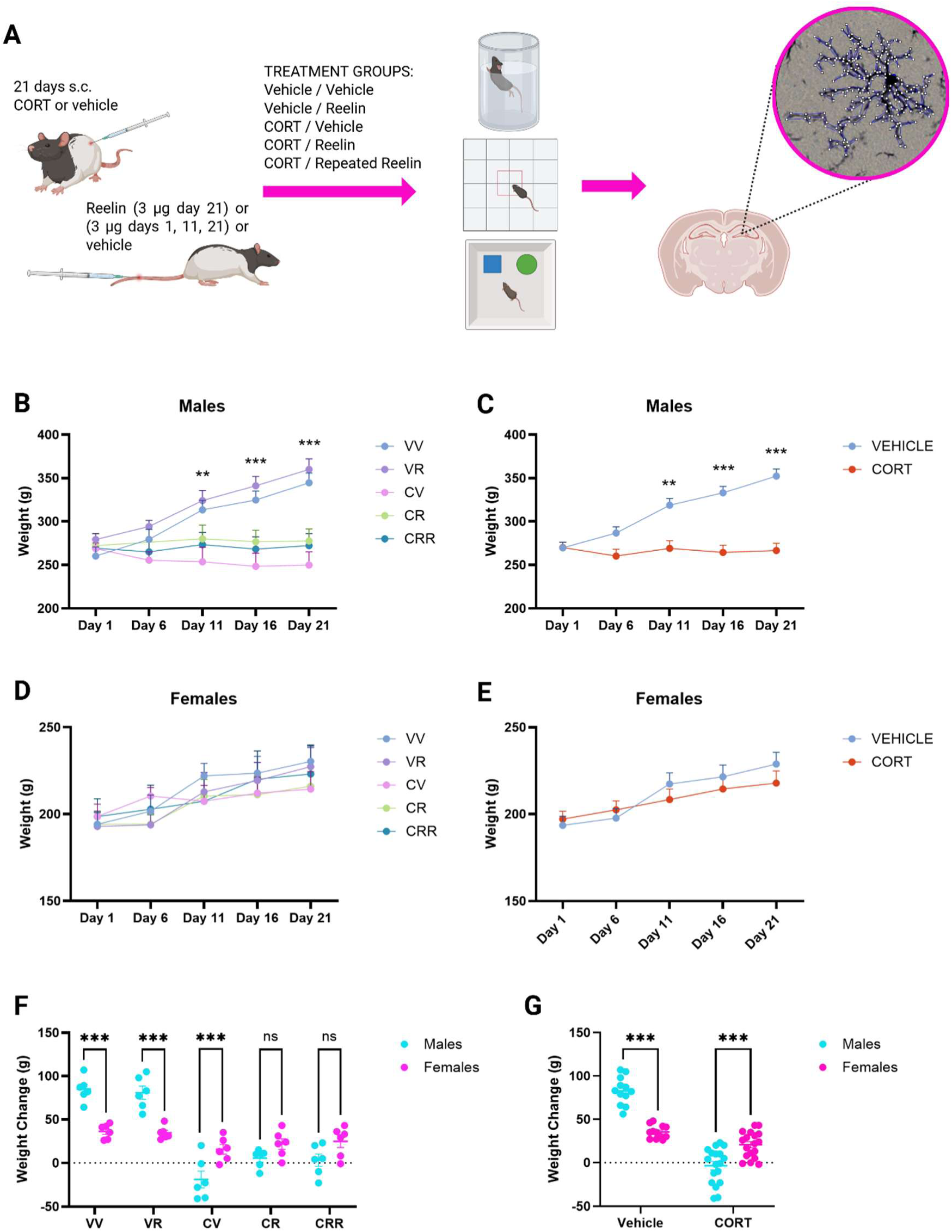
Animal model of chronic stress with single or multiple injections of reelin, and daily weights of rats. **(A)** Long Evans rats received daily subcutaneous injections of vehicle (0.9% NaCl with 2% polysorbate-80) or corticosterone (40 mg/ kg, suspended in vehicle). Reelin (3μg) or vehicle (0.1M phosphate-buffered saline, pH 7.4) was delivered via the lateral tail vein on days 1, 11, and 21. Rats underwent behavioural tests (forced swim test, open field test, and object location test) prior to perfusions with 4% paraformaldehyde. V=vehicle; C=CORT; R= Reelin. **(B-E)** Body weight for male and female rats over the 21-day injection protocol. A mixed model two-way ANOVA with stress and time as factors followed by Šídák’s multiple comparisons test shows that CORT significantly decreased weight gain for male rats (time *F*_(1.918,46.50)_ = 79.90, p < 0.001; CORT *F*_(1,28)_ = 18.85, p < 0.001; interaction *F*_(1.918,46.50)_ = 99.51, p < 0.001) but not female rats. **(F,G)** Change in body weight was higher for males receiving vehicle, but lower for males receiving CORT compared to female rats (sex *F*_(1,56)_ = 7.644, p = 0.08, CORT *F*_(1,56)_ = 143.7, p < 0.001), interaction *F*_(1,56)_ = 72.46, p < 0.001). Full statistical results are available in Supplementary Table S1. V=vehicle; C=CORT; R= Reelin. All data expressed as mean ± SEM; *p ≤0.05, **p ≤0.01, ***p ≤0.001.

### 3.2. Reelin rescued CORT-induced immobility in the forced swim test, but did not affect behaviour in the open field test or novel object location test

In the forced swim test, for both male and female rats, daily injections of CORT significantly increased immobility in the forced swim test (p < 0.001), which was reversed by both a single and multiple injections of reelin (Fig. 3A; males p = 0.003, p < 0.001; females p = 0.02, p = 0.006).

**Figure 3.**
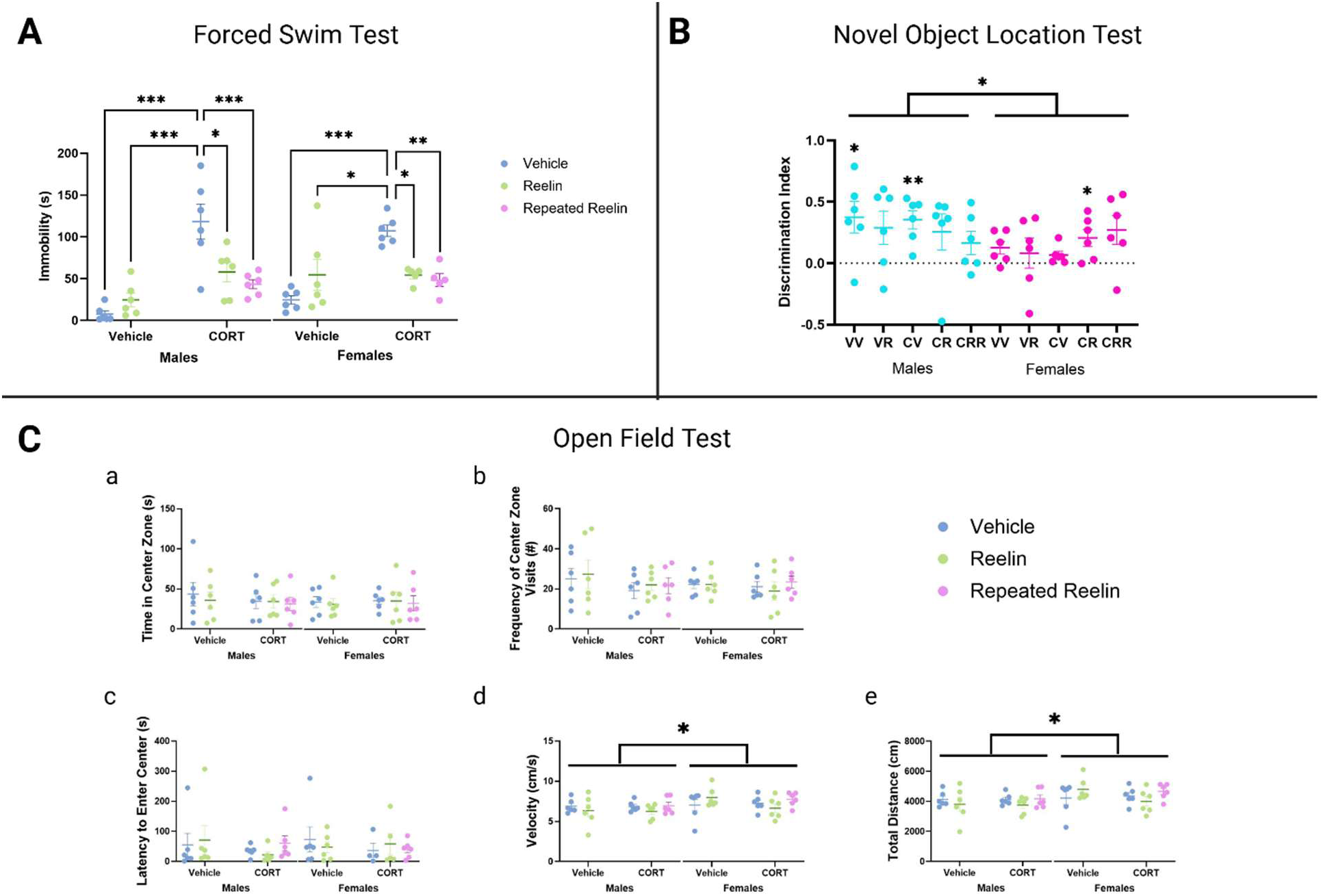
Behavioural results. **(A)** Forced swim test (FST). Rats were allowed to swim for 10 minutes and immobility was scored in the final 5 minutes of the test. A three-way ANOVA (CORT x reelin x sex) revealed significant main effects of CORT (*F*(1,48) = 38.75, p < 0.001) and reelin (*F*(2,48) = 10.15, p < 0.001), and a significant interaction between CORT and reelin (*F*(1,48) = 25.43, p < 0.001). Šídák’s multiple comparisons test showed that CORT administration significantly increased immobility in the FST for both male and female rats, which was decreased with multiple or a single injection of reelin. (**B**) Novel Object Location Test (NOLT). Rats were allowed to explore the objects placed in diagonal corners of the box for 4 minutes in the training phase. In the test phase, one object was moved to an adjacent corner. Rats explored the objects for 2 minutes, and time spent exploring both objects was scored. The discrimination index was calculated as: *(time with novel object – time with familiar object)/ (time with novel object + time with familiar object*). A three-way ANOVA (CORT x reelin x sex) indicated a significant main effect of sex (*F*_(1,48)_ = 4.310, p = 0.04). One sample t tests against a theoretical mean of 0 showed that male VV rats, male CV rats, and female CR rats spent significantly more time with the displaced object. (C) Open field test (OFT). Rats were left to explore an OFT box for 10 minutes. Data were analyzed using a three-way ANOVA (CORT x reelin x sex), followed by Šídák’s post hoc multiple comparisons test. CORT did not produce anxiety-like behaviour in either sex as evaluated by time in the center zone, frequency of center zone visits, latency to enter the center zone, velocity, and total distance travelled. Overall, females were more mobile than males (sex *F*_(1,48)_ = 4.81, p = 0.03). Full statistical results are available in Supplementary Table S1. V=vehicle; C=CORT; R= Reelin. All data is presented as mean ± SEM; *p < 0.05, **p ≤ 0.01, ***p ≤ 0.001.

In the novel object location test, the DI describes the percentage of time spent exploring the object that had moved (DI 0% to 100%; expressed as 0 to 1), and the object that had not moved (DI 0% to −100%; expressed as 0 to −1) (Fig. 3B). A DI of 0% indicates equal exploration of both objects. The DI is calculated as follows: (time spent exploring the displaced object – time spent exploring the familiar object)/ (*time spent exploring the displaced object + time spent exploring the familiar object*). Measurements were compared to DI = 0 to determine significance. Male rats that received vehicle only differed from chance by 0.37, 95% CI [0.04, 0.70], as with male rats that had received CORT only (0.35, 95% CI [0.16, 0.54]). For female rats, only rats that received one dose of reelin following daily CORT differed from chance (0.21, 95% CI [0.02, 0.38]).

In the open field test, females were significantly more mobile than males, as shown by increased velocity and distance travelled (Fig. 3C; p = 0.03), however no significant effects of treatment were found.

### 3.3. Microglia density did not change with treatment in the polymorphic layer or subgranular zone of the dorsal hippocampus

A three-way ANOVA showed a significant main effect of CORT in the PML (*F*_(1,48)_ = 6.60, p = 0.01). No significant group comparisons were revealed with Šídák’s post-hoc comparison test in either the PML or the SGZ. Density results are shown in Figure 4.

**Figure 4.**
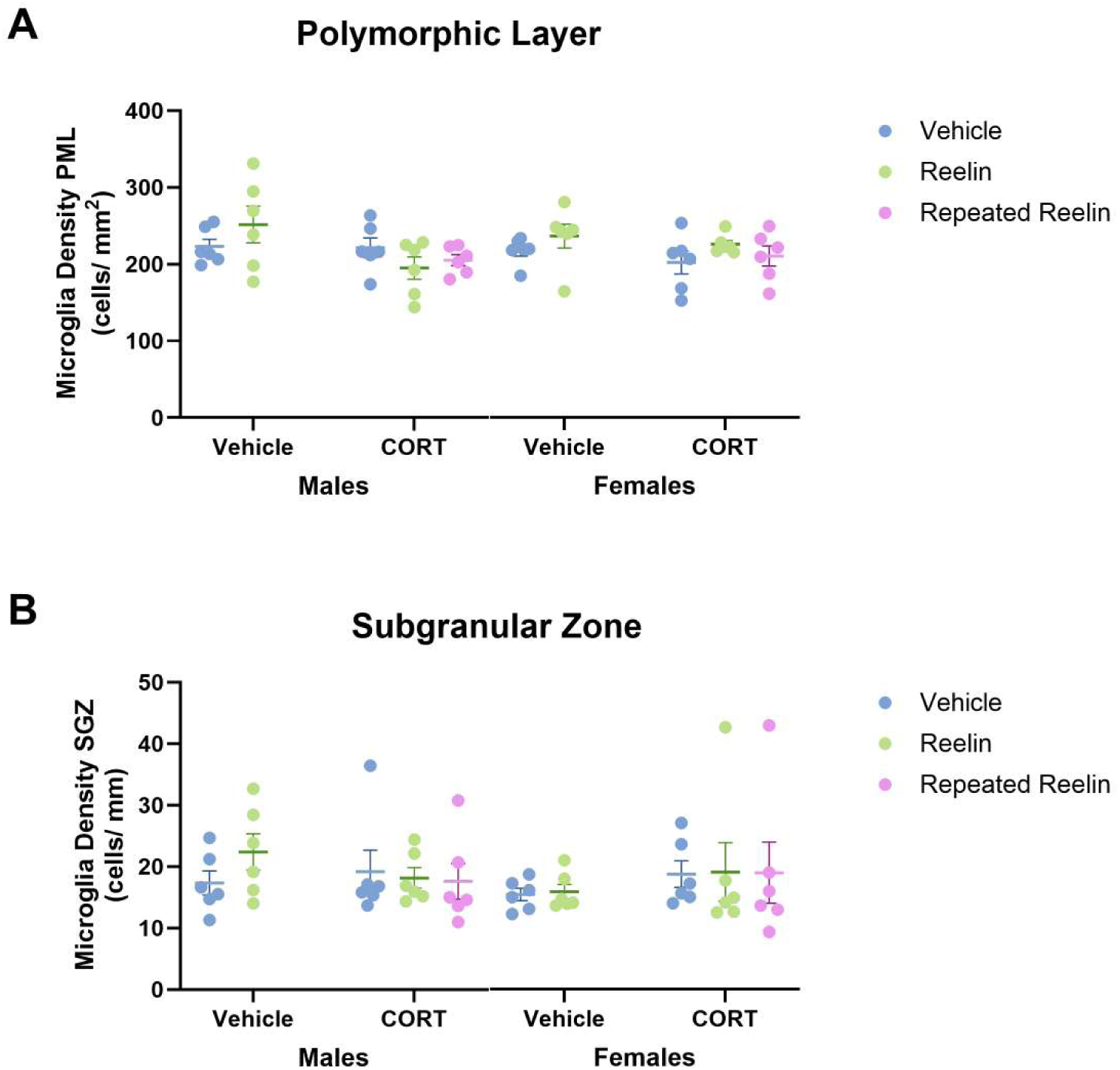
Microglia density in the polymorphic layer (PML) and subgranular zone (SGZ) of the dorsal hippocampus. **(A)** Microglia density across treatment groups did not significantly differ in the PML, although there was a significant main effect of CORT (*F*(1,48) = 6.60, p = 0.01). **(B)** No significant effect of treatment of sex was found in the SGZ. Statistical analyses were carried out using a three-way ANOVA (CORT x reelin x sex) with Šídák’s multiple comparison post hoc test. Full statistical results are available in Supplementary Table S1. V=vehicle; C=CORT; R= Reelin. All data is presented as mean ± SEM; *p < 0.05, **p ≤ 0.01, ***p ≤ 0.001.

### 3.4. Chronic CORT and reelin produced opposing effects on microglia morphological parameters – process tracings

Microglia process tracing results suggest that 21 days of CORT reduced branching of processes, and reelin treatment alone (V/R group) increased branching. CORT reduced total process length (*F_(_*_1,48)_ = 4.16, p = 0.05) and total intersections (F_(1,48)_ = 5.60, p = 0.02) regardless of reelin treatment, and reduced all measured parameters at multiple distances from the soma (Fig. 5). There was a significant difference in V/R and C/V microglia morphology on multiple measured parameters, suggesting that CORT and reelin may have opposing effects on microglia morphology. Chronically stressed rats had reduced nodes at more distal radii (Fig. 5C,G). Additionally, there was a main effect of sex in multiple parameters across numerous distances from the soma, with an increased number of significant alterations in male rats. Full

**Figure 5.**
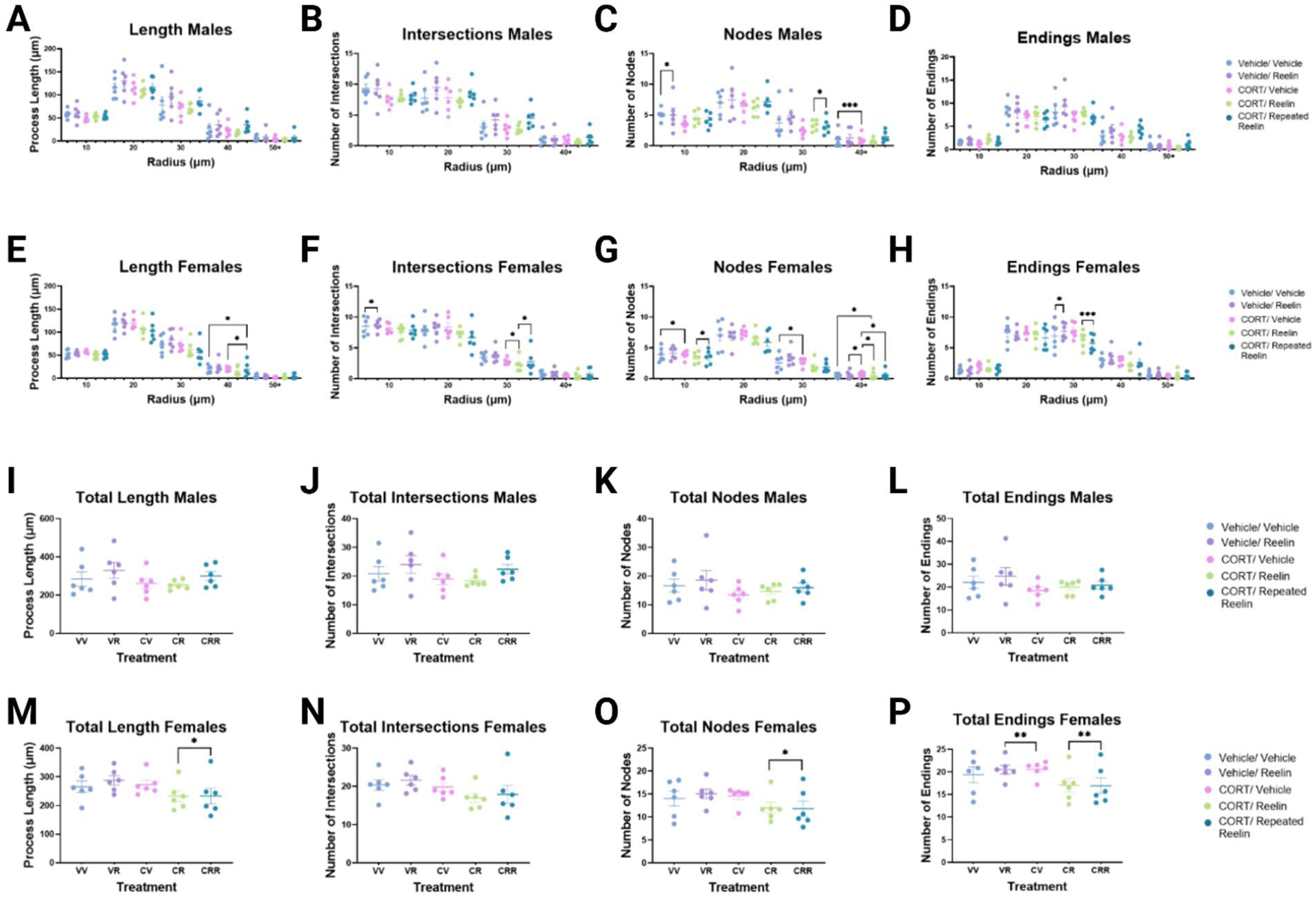
Microglia process tracings in the polymorphic layer of the hippocampus. Processes of six microglia per rat were traced manually in Neurolucida. Morphology was determined for process length **(A,E,I,M),** process intersections **(B,F,J,N),** process nodes **(C,G,K,O)**, and process endings **(D,H,L,P**) using a Sholl analysis with radii at 10, 20, 30, 40, and 50 *μ*m from the soma. Data were analyzed in R using linear mixed-effects models with CORT, Reelin, and Sex as fixed factors, followed by Šídák-adjusted post hoc comparisons. Full statistical results are available in Supplementary Table S1. V=vehicle; C=CORT; R= Reelin. All data presented as mean ± SEM; *p ≤0.05, **p ≤0.01, ***p ≤0.001.

### 3.5. Microglia outline tracings showed stress altered microglia morphology, which was reversed with single or repeated Reelin injections

#### 3.5.1. Polymorphic layer dentate gyrus

In both male and female rats, chronic CORT significantly decreased microglia area (males p < 0.001; females p = 0.006; Fig. 6A), perimeter (males p < 0.001; females p < 0.001; Fig. 6B), and max feret (males p <0.001; females p <0.001; Fig. 6D), and increased circularity (males p < 0.001; females p = 0.004; Fig. 6C), aspect ratio (males p = 0.02; females p = 0.007; Fig. 6E), and roundness (females p = 0.04; Fig. 6F). CORT alone did not alter solidity for either sex as determined with Šídák’s multiple comparisons test (Fig. 6G), but there were main effects of CORT (*F*_(1,48)_ = 29.08, p < 0.001) and reelin (*F*_(2,48)_ = 13.21, p < 0.001).

**Figure 6.**
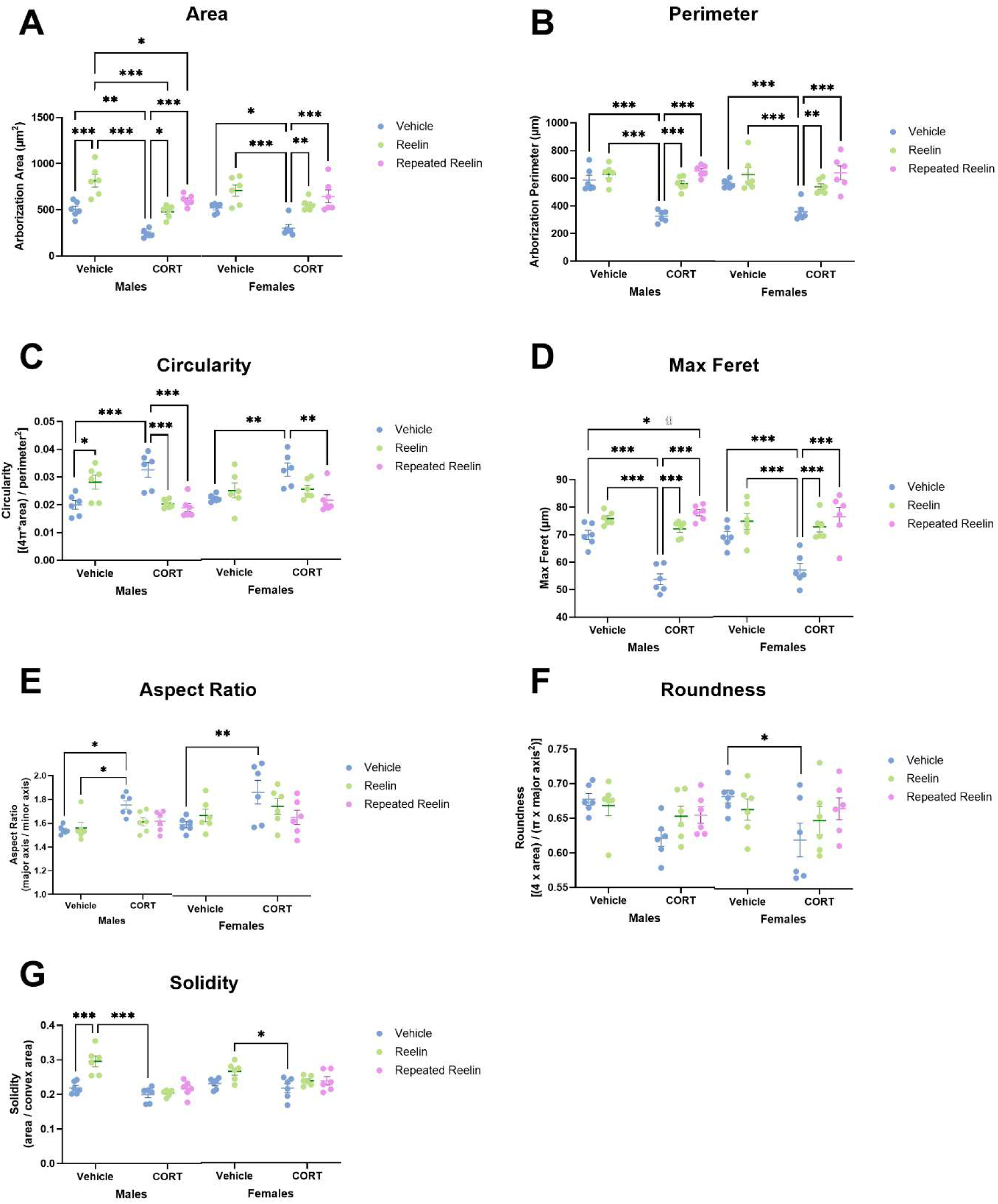
Reelin rescued CORT-induced alterations to polymorphic layer microglia morphology in male and female rats. Outlines of 25 microglia per rat in the polymorphic layer of the dentate gyrus were manually traced in ImageJ. Microglia morphology was assessed based on the following morphological parameters: cell area **(A),** perimeter **(B),** circularity (ranges from 0-1, with 1 being a perfect circle) **(C),** max feret (the longest distance between any two points on the cell’s perimeter) **(D),** aspect ratio (1 is a perfect circle, >1 is more elongated and less symmetrical) **(E),** roundness **(F)**, and solidity (close to 1 is a smooth, compact outline; <1 indicates irregularity in the perimeter with many branches or indentations) **(G).** CORT produced process retraction and smaller cells, which was normalized with one or multiple injections of reelin for the majority of measured parameters. Data were analyzed in R using linear mixed-effects models with CORT, reelin, and sex as fixed factors, followed by Šídák-adjusted post hoc comparisons. Full statistical results are available in Supplementary Table S1. V=vehicle; C=CORT; R= Reelin. All data presented as mean ± SEM; *p ≤0.05, **p ≤0.01, ***p ≤0.001.

Taken together, these parameters suggest that CORT produces smaller cells (↓ area, ↓ feret) with retracted processes (↓ perimeter, ↓ feret) and a more symmetrical perimeter (↑ aspect ratio, ↑ circularity). Reelin alone (V/R group) showed the opposite effect of CORT for most parameters, increasing cell size (↑ area, Fig. 6A) for males (p < 0.001) and females (p = 0.03), and for males only, process bushiness (↑ solidity; p < 0.001; Fig. 6G).

Reelin administration following chronic CORT effectively normalized morphology across the measured parameters. For males and females, both a single injection (3 µg total) and repeated injections (9 µg total) of reelin reversed CORT-induced decreases in area (males p = 0.003, females p < 0.001) and perimeter (males p < 0.001, females p < 0.001); Fig. 6A, B), and CORT-induced increases in feret (males p < 0.001, females p < 0.001; Fig. 6D). While both doses of reelin after chronic CORT also reversed increased circularity for males (p < 0.001, p < 0.001; Fig. 6C), only repeated reelin was sufficient to significantly decrease circularity in females (p = 0.002; Fig. 6C). Aspect ratio was not significantly different from V/V or from C/V groups following a single injection or repeated injections of reelin for both sexes (Fig. 6E).

#### 3.5.2. Subgranular zone dentate gyrus

Contrarily to morphological alterations observed in the PML, microglia in the SGZ did not show significantly altered morphology following chronic stress or reelin treatments as determined by Šídák’s multiple comparisons test (Fig. 7). However, significant interaction effects for CORT:Reelin:Sex were found for area (*F*_(1,48)_ = 4.77, p = 0.03) and circularity (*F*_(1,48)_ = 7.70, p = 0.008), and for CORT:Sex for circularity (*F*(1,48) = 4.93, p = 0.03). Circularity was elevated following chronic stress for males and decreased with reelin treatment, suggesting a possible similar effect of treatment, although not significant in this region.

**Figure 7.**
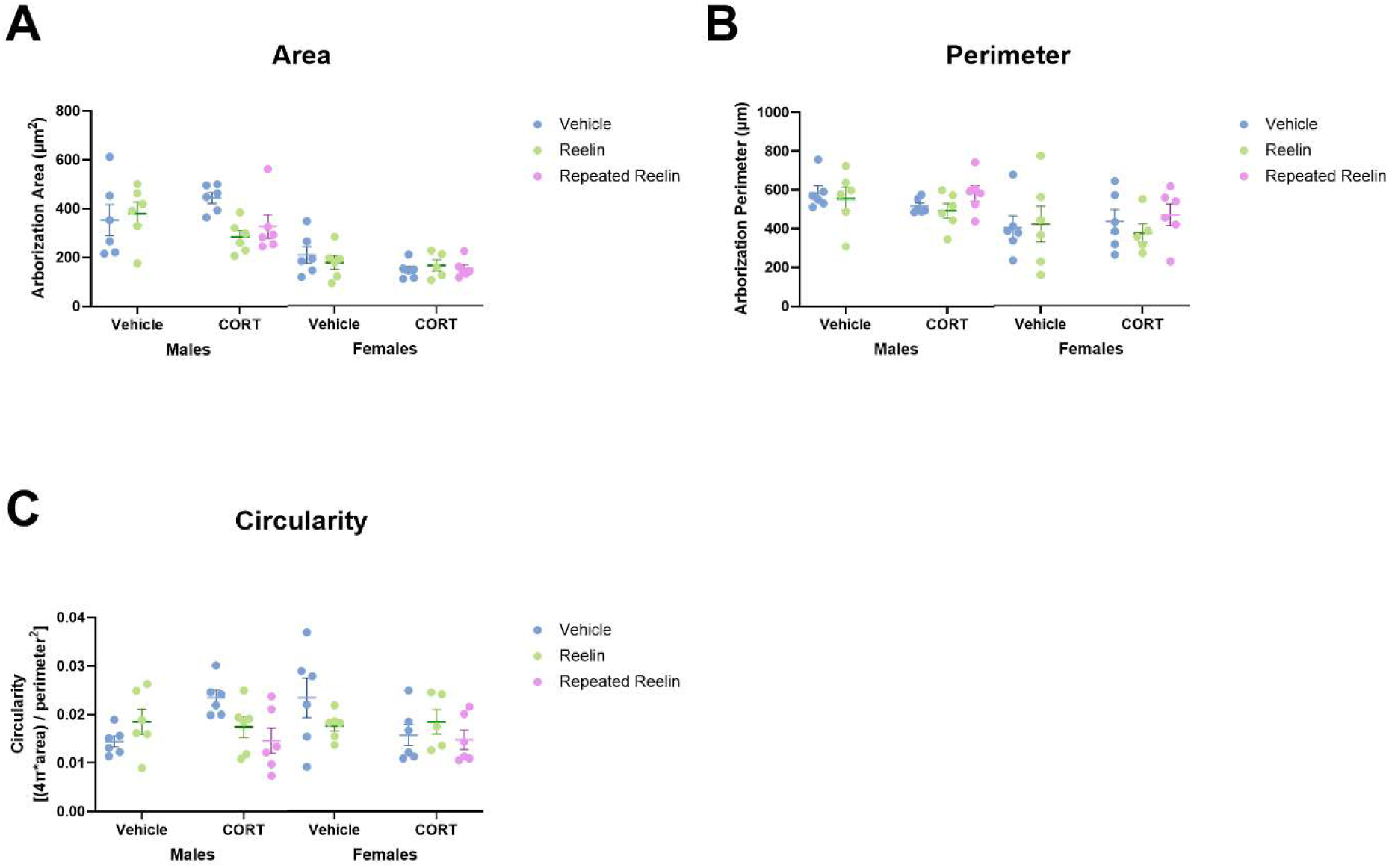
Microglia morphology in the subgranular zone did not significantly change with treatment. 25 microglia per rat were traced using QuPath in the subgranular zone of the hippocampal dentate gyrus. Measure of area, perimeter, and circularity were recorded. Data were analyzed in R using linear mixed-effects models with CORT, reelin, and sex as fixed factors, followed by Šídák-adjusted post hoc comparisons. Full statistical results are available in Supplementary Table S1. V=vehicle; C=CORT; R= Reelin. All data presented as mean ± SEM.

### 3.6. CORT and/ or reelin did not change microglia soma area

Soma areas of microglia did not differ between treatment groups for either males or females (Fig. 8), and no significant main effect of treatment or sex was observed with a nested linear mixed effects model.

**Figure 8.**
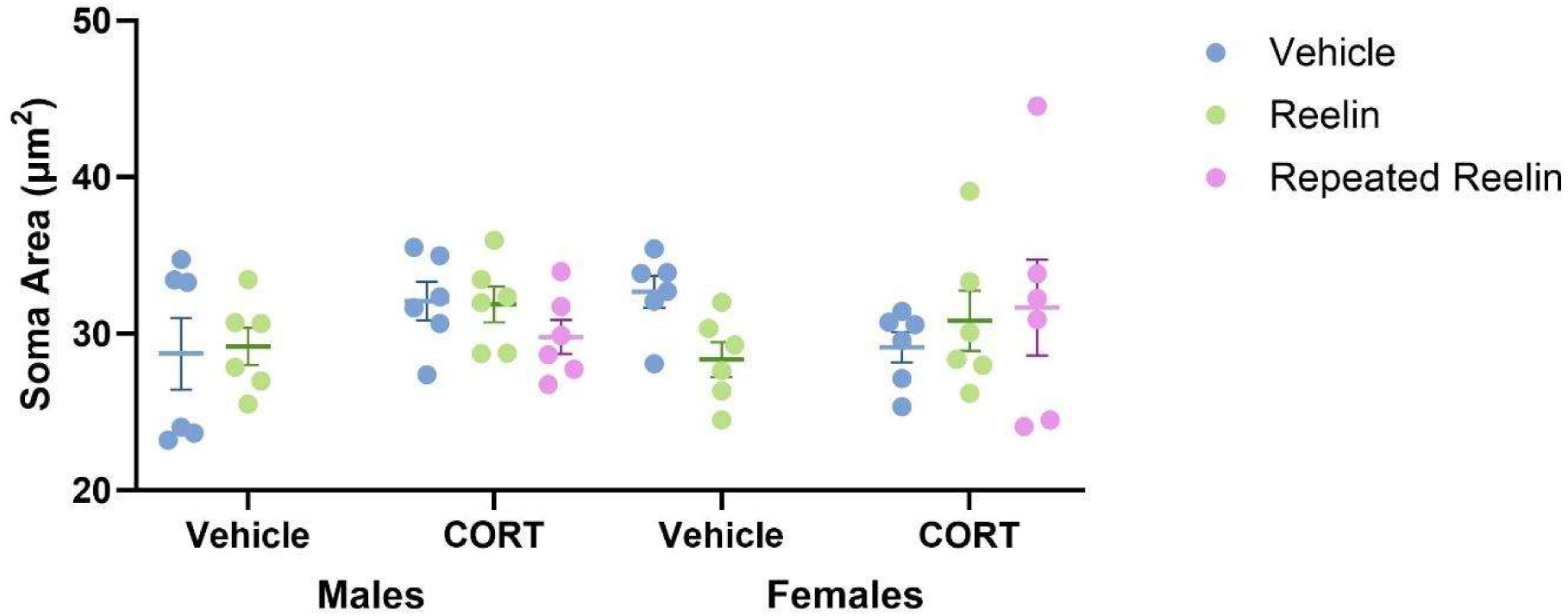
Soma area did not differ following daily injections of CORT or after Reelin treatment. Somas from 50 microglia per rat were traced in the polymorphic layer (PML) of the hippocampus with ImageJ, and soma area was measured. Data were analyzed in R using linear mixed-effects models with CORT, reelin, and sex as fixed factors. No significant effects of treatment were found. Full statistical results are available in Supplementary Table S1. V=vehicle; C=CORT; R= Reelin. All data presented as mean ± SEM.

### 3.7. CORT increases % of microglia expressing cleaved caspase-3 for female rats, which is normalized with reelin

In the PML, the C/V group did not differ significantly from the V/V group, but both a single and multiple injections of reelin decreased the percentage of Iba1-IR microglia expressing cleaved caspase-3 as compared to CORT only for female rats (p = 0.003; p = 0.001). In the SGZ, CORT administration increased the percentage of Iba1-IR microglia expressing cleaved caspase-3 (p = 0.01), which was normalized with both single (p = 0.03) and multiple injections of reelin (p = 0.002). Cleaved caspase-3 expression is shown in Figure 9.

**Figure 9.**
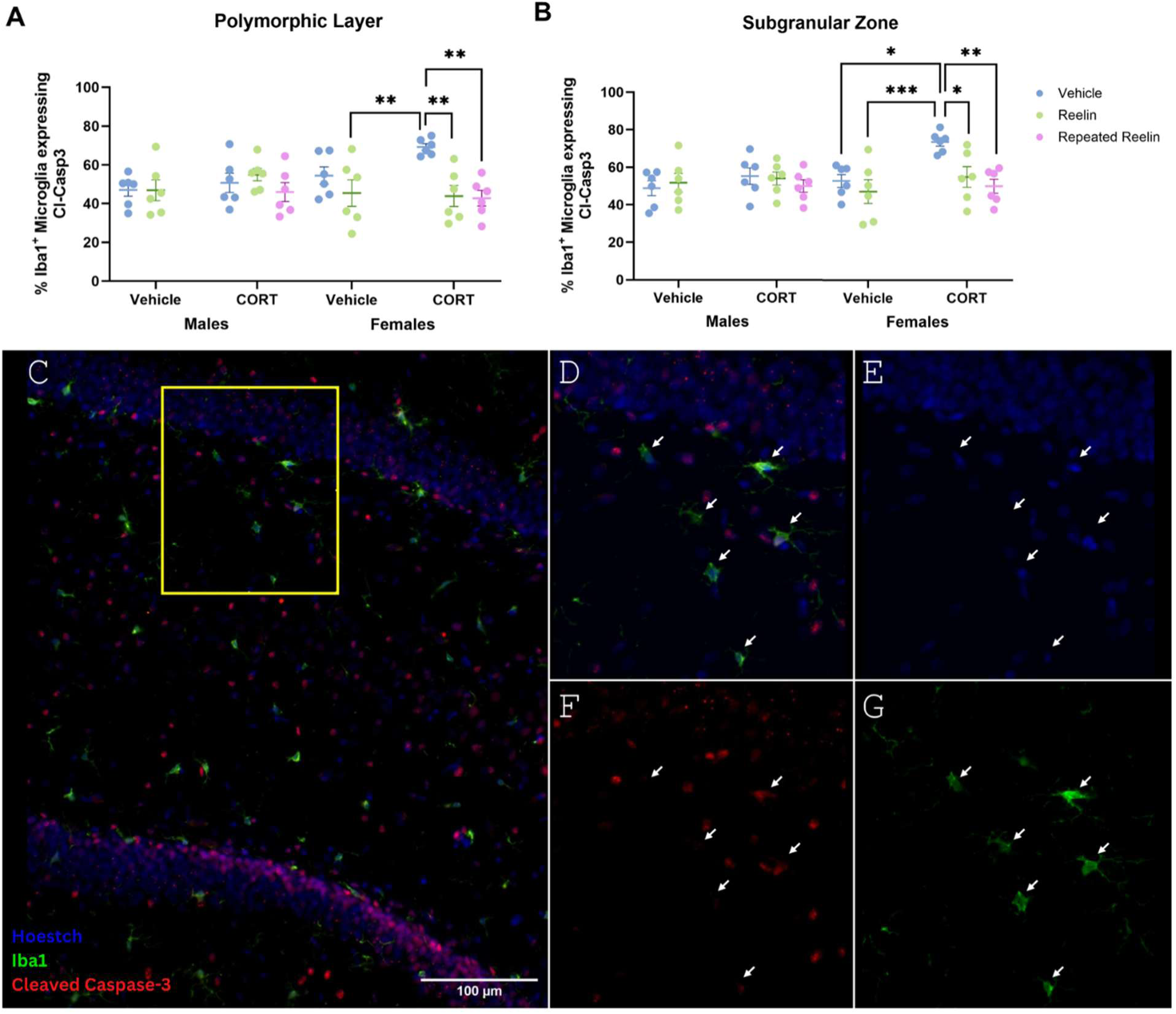
CORT increases cleaved caspase 3 (cleaved caspase-3) expression in hippocampal microglia in female rats. **(A)** polymorphic layer, **(B)** subgranular zone. Sections containing the dorsal hippocampus were stained for Iba1 and cleaved caspase-3 to assess cleaved caspase-3 expression in microglia. Cleaved caspase-3 expression was determined in at least 100 microglia per rat, and the proportion of microglia expressing cleaved caspase-3 (Iba1^+^cl-cas3^+^ / Iba1^+^cleaved caspase-3^-^) was expressed as a percentage. A three-way ANOVA (CORT x reelin x sex) with Šídák’s multiple comparisons post hoc test revealed that 21 days of CORT increased the percentage of hippocampal microglia expressing cleaved caspase-3 in female rats, which was normalized with reelin administration. No effect of treatment was observed in male rats. **(C)** Iba1^+^ microglia in the dorsal hippocampus (20x). **(D-E)** Hoestch nucleus stain (blue); cleaved caspase-3 (red); Iba1^+^ microglia (green). Full statistical results are available in Supplementary Table S1. V=vehicle; C=CORT; R= Reelin. All data presented as mean ± SEM; *p ≤0.05, **p ≤0.01, ***p ≤0.001.

## 4. Discussion

In this study, we evaluated if intravenous reelin promotes microglia morphologies suggestive of neuroinflammation following therapeutic and escalated dosage schedules. We show that reelin normalizes chronic stress-induced alterations to hippocampal microglia morphology when delivered at a dose found to be effective in reversing stress-associated behavioural deficits, and further show that multiple injections of reelin may bring about pro-inflammatory changes. We additionally show a female-specific increase in microglia IR for cleaved caspase-3 following chronic CORT that is resolved with reelin administration. These results suggest an acute dose of 3 µg of reelin, which reverses both behavioural and neurobiological effects of chronic stress, also reverses morphological and possibly functional changes to hippocampal microglia induced by chronic stress and does not lead to effects consistent with the enhancement of pro-inflammatory effects (Allen et al., 2022; Johnston et al., 2023; Reive et al., 2026). We also observed sex differences in the impact of reelin on hippocampal microglia morphology. Altogether these results suggest reelin homeostasis is a valuable and targetable pathway in the contexts of both depression and inflammation. This, along with past research showing that reelin normalizes numerous chronic stress-induced behavioural and physiological alterations, corroborates the possibility of developing reelin-based therapeutics for major depression (Brymer et al., 2020; Johnston et al., 2023; Reive et al., 2023; Sánchez-Lafuente et al., 2024).

Male and female rats were exposed to chronic stress and received either a single injection of reelin on the final day of chronic stress, or repeated (3) reelin injections throughout chronic stress. Behaviourally, reelin significantly reduced CORT-induced increases in immobility in the FST within 24 hours of administration (we show here 6 hours post injection) as seen in past studies (Allen et al., 2022; Johnston et al., 2023; Reive et al., 2023; Scheil et al., 2024). This was true for rats treated repeatedly with reelin and rats receiving a single injection. No difference to locomotion was observed in the open field test following either chronic stress or reelin administration, indicating the antidepressant-like effect of reelin as measured by reduced immobility in the FST is not attributable to non-specific psychostimulant or locomotor effects. CORT induced process retraction, and both a single and repeated intravenous injections of reelin reversed changes to microglia morphology in male and female rats. Additionally, a single injection of reelin in vehicle rats increased the solidity, circularity, and area of the traces without altering perimeter, suggesting that elevated levels of reelin may induce thickened processes consistent with a shift towards primed microglia. Alternatively, microglia from rats exposed to chronic CORT or reelin alone may be transitioning towards an amoeboid-like morphology commonly observed in disease states; however, amoeboid-like morphologies are marked by an enlarged soma which was not found here. This is consistent with our previous investigation of the effects of chronic CORT and reelin on microglia morphology in which no significant differences to soma size were found (Reive et al., 2026).

We show that a dose of reelin, which rapidly reversed chronic CORT-induced despair-like behaviour, leads to a normalization of stress-induced morphological changes in hippocampal microglia, and that reelin administration in non-stressed rats fosters a hyper-ramified morphology previously associated with a transition state towards reactive microglia as well as disease states including chronic stress (Hellwig et al., 2016; Hinwood et al., 2013; Reddaway et al., 2023; Rowson et al., 2016). These results support our recent observations of reelin impacting microglia morphology, where we found multiple injections of reelin increased process complexity in CORT and vehicle-treated male rats (Reive et al., 2026). Elevating reelin levels through multiple peripheral reelin injections causing pro-inflammatory-like microglial changes to microglia morphology is consistent with past studies suggesting reelin has pro-inflammatory effects (Alexander et al., 2023; Calvier et al., 2020; Reive et al., 2024). Microglia have been shown to express reelin’s canonical receptor VLDLR, and therefore may interact directly with reelin but changes to peripheral inflammatory signalling could also be responsible for alterations to microglia morphology following reelin injection(s) (Fan et al., 2001; Pocivavsek et al., 2009). Additionally, microglia respond to changes in the extracellular matrix, and dysregulation of extracellular matrix proteins such as reelin could also influence microglia morphology and functional states (Wareham & Calkins, 2025). As reelin signals through VLDLR and extracellular signal-regulated kinase (ERK), reelin in the CNS may bind to microglia to influence microglia-invoked inflammation or other microglia functions. Reelin could also influence process extension and branching through promoting actin polymerization and microtubule stabilization, paralleling its effects in neurons to promote cytoskeleton reorganization and dendritogenesis (Bórquez et al., 2013; Chai et al., 2009). Reelin may alternatively affect microglia morphology indirectly through promotion of hippocampal long-term potentiation (LTP) (Johnston et al., 2023). Reelin administration following chronic stress increases LTP in the hippocampus, and enhancement of LTP triggers microglia process formation, branching, and increased microglia contact with dendritic spines (Johnston et al., 2023; Pfeiffer et al., 2016). The effects of reelin on microglia morphology can be direct or indirect, and although we report morphological changes to hippocampal microglia in this study, the responsible mechanisms remain unknown and require further evaluation.

We found no effect of treatment on microglia density in either the PML or SGZ following cell counts of Iba1-IR microglia. Previous studies show microglia proliferation in the hippocampus following shorter periods of chronic stress (Tynan et al., 2010), but longer-term chronic stress protocols decrease microglia density (Kreisel et al., 2014; Tong et al., 2017). It is possible that we did not observe changes in microglia density as our 3-week chronic stress paradigm falls between the durations typically associated with proliferation (days to 2 weeks; (Chen et al., 2025; Lehmann et al., 2016; Nair & Bonneau, 2006; Tynan et al., 2010)) and microglia loss (4-5 weeks; (Kreisel et al., 2014; Tong et al., 2017)).

We observed no noteworthy sex differences when considering hippocampal microglia morphology; however, we did observe a female-specific increase in cleaved caspase-3 immunoreactivity. Evidence from both clinical and preclinical studies indicates distinct sex-specific neuroimmune responses to chronic stress (Bekhbat & Neigh, 2018). Women are diagnosed with depression almost twice as often as men, and appear more vulnerable to the mood and social depressogenic effects of inflammation, despite a heterogeneous pattern of peripheral biomarker associations with depression across sexes (Albert, 2015; Eisenberger et al., 2009; Kropp & Hodes, 2023; Moieni et al., 2015). Female rats display a significantly higher number of microglia with an activated phenotype than males in brain regions associated with the pathogenesis of depression, including the hippocampus (Schwarz et al., 2012). Female mice have been found to have heightened baseline and stress-induced CORT levels, indicating a potential for an exaggerated response to chronic stress and a higher susceptibility to neuroinflammatory priming (Frank et al., 2014; Solomon et al., 2015; Tinnikov, 1999). Additionally, some evidence shows elevated hippocampal interleukin (IL)-6 levels in female mice but not male mice following chronic stress and an acute LPS injection, suggesting a sex-dependent and hippocampus-dependent effect (Careaga & Wu, 2024). Conversely, an ex vivo study reports elevated microglia-specific cytokine responses in microglia from stressed male rats only, highlighting contrasting findings in comparing male and female neuroimmune responses in vivo and ex vivo (Fonken et al., 2018). We evaluated microglial cleaved caspase-3 immunoreactivity as an additional indicator as to whether peripheral reelin injections impact CNS inflammation. In female rats only, the proportion of Iba1-stained microglia co-expressing cleaved caspase-3 was increased with repeated CORT injections, and normalized with a single or repeated reelin injection(s). Although mechanistic insights of reelin’s influence in caspase-3 signalling in microglia is quite limited, a female-specific elevation in cleaved caspase-3 is consistent with aforementioned research indicating an exaggerated immune response to chronic stress in females compared to males (Careaga & Wu, 2024; Frank et al., 2014; Solomon et al., 2015). Furthermore, apolipoprotein E signalling through microglial VLDLR increases extracellular signal-regulated kinase (ERK) activation and decreases c-Jun N-terminal kinase (JNK) activation to inhibit a lipopolysaccharide (LPS)-induced inflammatory response (nitric oxide production), suggesting that reelin may similarly attenuate microglia-invoked inflammation (Pocivavsek et al., 2009).

Cleaved caspase-3 signalling differentially regulates cell death and inflammatory activation (Burguillos et al., 2011; Kavanagh et al., 2014; Shen et al., 2016). In response to an inflammatory stimulus, upstream caspases cleave pro-caspase-3, generating caspase-3 p19/12 complexes. These caspase-3 p19/12 complexes can activate cytoplasmic nuclear factor kappa B (NF-κB) through protein kinase C (PKC), which translocates to the nucleus to regulate the transcription of immune-related genes (Kavanagh et al., 2014). Caspase-3 p19/12 can also form p17/12 complexes following autocatalytic processing, which translocate to the nucleus to induce cell death. There is evidence for a distinct cleaved caspase-3 expression profile, with caspase-3 p19 expression restricted to the cytoplasm, while caspase-3 p17 expression present in both the cytoplasm and nuclear fractions (Kavanagh et al., 2014). As we observed cleaved caspase-3 staining in the cytoplasm as well as the nucleus, it is plausible that increased cleaved caspase-3 expression in female rats exposed to 21 days of CORT injections is indicative of an inflammatory response involving microglial caspase-3 rather than apoptosis, especially considering no changes in microglia density were found. Although further assessments are required to determine the responsible mechanisms, female-specific increases in cleaved caspase-3 found in this study could be an indicator of an exaggerated neuroimmune response to chronic CORT. It will be valuable to determine whether analysis of hippocampal cytokines corroborates our observations following exposure to chronic CORT and reelin in future studies.

While the present results establish a novel insight into microglia dynamics following exposure to reelin, certain limitations that should be acknowledged. Although morphological changes often parallel inflammatory activation and can serve as an indirect indicator of neuroimmune status, morphology alone is insufficient to fully capture the complexity of inflammatory dynamics (Paolicelli et al., 2022). Additionally, while we did observe microglia that were IR for Iba1 but not cleaved caspase-3, the strong Iba1 signal may have resulted in channel bleed-through, potentially contributing to inflated estimates of Iba1/cleaved caspase-3 colocalization. Finally, we delivered daily CORT, a potent anti-inflammatory compound, at supraphysiological levels to create a phenotype of chronic stress. Immune cells can develop resistance to the anti-inflammatory effects of glucocorticoids following prolonged exposure or exposure to high doses, driving pro-inflammatory processes. Therefore, the present results do not definitively establish whether reelin acts specifically as an anti-inflammatory compound; however, they do demonstrate that reelin counteracted many of the effects of prolonged CORT exposure.

## CONCLUSION

Taken together, these findings indicate that a dose of reelin reversing behavioural and neuroplasticity consequences of chronic stress does not produce neuroinflammatory consequences following chronic stress as characterized by microglia morphology and cleaved caspase-3 expression. Peripheral reelin injections can modify inflammatory states in the CNS, which carries significant implications for neuropsychiatric conditions marked by inflammation and impaired reelin homeostasis. Additionally, we demonstrated therapeutic doses of reelin can reverse several neuroinflammatory consequences of chronic stress in male and female rats as determined through assessment of microglia morphology and cleaved caspase-3 expression. These results support reelin homeostasis as an important consideration in the treatment of depression and strengthen the rationale for continued advancement of reelin towards clinical application.

## Funding statement

This work was supported by the CIHR CGRS-D to CSH (Funding Reference Number 210219), NSERC DG to HJC and LEK, and CIHR PG and CRC to HJC.

## Notes

### Competing Interest Statement

The authors have declared no competing interest.

