## Supplementary Table S1 for "Reelin doses normalizing behavioural alterations induced by repeated corticosterone avoid eliciting neuroinflammation-associated morphology in hippocampal microglia"

| Figure Number | Figure Description | Statistical Test | n | p value | F values | Partial Eta Squared | R Squared | Post-Hocs | Post-Hocs Females | Post-Hocs Males | Cohen's d Fem | Cohen's d Mal | Westfall d Females | Westfall d Males |
|---|---|---|---|---|---|---|---|---|---|---|---|---|---|---|
| 2B-C | Body weight for male rats over the 21-day injection protocol | Two-way ANOVA Greenhouse–Geisser correction<br>Post hoc:<br>Šídák's multiple comparisons test | Vehicle/Vehicle = 6<br>Vehicle/Reelin = 6<br>CORT/Vehicle = 6<br>CORT/Reelin = 6<br>CORT/Repeated Reelin = 6 | Interaction: p < 0.001<br>CORT: p < 0.001<br>Time: p < 0.001 | F(1.918,46.50) = 99.51<br>F(1,28) = 18.85<br>F(1.918,46.50) = 79.90 | | | | | Day 1: p = 1<br>Day 6: p = 1.01<br>Day 11: p = 0.02<br>Day 16: p < 0.001<br>Day 21: p < 0.001 | | | | |
| 2D-E | Body weight for female rats over the 21-day injection protocol | Two-way ANOVA<br>Post hoc:<br>Šídák's multiple comparisons test | Vehicle/Vehicle = 6<br>Vehicle/Reelin = 6<br>CORT/Vehicle = 6<br>CORT/Reelin = 6<br>CORT/Repeated Reelin = 6 | Interaction: p < 0.001<br>CORT: p = 0.735<br>Time: p = 0.002 | F(2.178, 55.55) = 95.16<br>F(1,28) = 0.12<br>F(2.178, 55.55) = 6.84 | | | | Day 1: p = 0.99<br>Day 6: p = 0.98<br>Day 11: p = 0.85<br>Day 16: p = 0.96<br>Day 21: p = 0.79 | | | | | |
| 2F | Change in body weight by sex and treatment group | Two-way ANOVA<br>Post hoc:<br>Šídák's multiple comparisons test | Vehicle/Vehicle = 6<br>Vehicle/Reelin = 6<br>CORT/Vehicle = 6<br>CORT/Reelin = 6<br>CORT/Repeated Reelin = 6 | Interaction: p < 0.001<br>Condition: p < 0.001<br>Sex: p = 0.27 | F(4,50) = 20.09<br>F(4,50) = 40.78<br>F(1,50) = 1.27 | | | VV: p < 0.001<br>VR: p < 0.001<br>CV: p = 0.001<br>CR: p = 0.30<br>CRR: p = 0.09 | | | | | | |
| 2G | Change in body weight by sex and CORT treatment | Two-way ANOVA<br>Post hoc:<br>Šídák's multiple comparisons test | Vehicle/Vehicle = 6<br>Vehicle/Reelin = 6<br>CORT/Vehicle = 6<br>CORT/Reelin = 6<br>CORT/Repeated Reelin = 6 | Interaction: p < 0.001<br>CORT: p < 0.001<br>Sex: p = 0.08 | F(1,56) = 72.46<br>F(1,56) = 143.7<br>F(1,56) = 7.644 | | | Vehicle (males) - Vehicle (females): p < 0.001<br>CORT (males) - CORT (females): p < 0.001 | | | | | | |
| 3A | Behavioural results - Forced swim test | Three-way ANOVA<br>Post hoc:<br>Šídák's multiple comparisons test | Vehicle/Vehicle = 6<br>Vehicle/Reelin = 6<br>CORT/Vehicle = 6<br>CORT/Reelin = 6<br>CORT/Repeated Reelin = 6 | CORT: p < 0.001<br>Reelin: p < 0.001<br>Sex: p = 0.27<br>CORT:Reelin: p < 0.001<br>CORT:Sex: p = 0.07<br>Reelin:Sex: p = 0.67<br>CORT:Reelin:Sex: p = 0.86 | F(1,48) = 38.75<br>F(2,48) = 10.15<br>F(1,48) = 1.23<br>F(1,48) = 25.43<br>F(1,48) = 3.53<br>F(2,48) = 0.40<br>F(1,48) = 0.03 | 0.446681911<br>0.297269945<br>0.024900999<br>0.34631322<br>0.068589549<br>0.016303589<br>0.000669251 | | | Vehicle/Vehicle - Vehicle/Reelin: p = 0.40<br>Vehicle/Vehicle - CORT/Vehicle: p < 0.001<br>Vehicle/Vehicle - CORT/Reelin: p = 0.48<br>Vehicle/Vehicle - CORT/Repeated Reelin: p = 0.75<br>CORT/Vehicle - Vehicle/Reelin: p = 0.01<br>CORT/Vehicle - CORT/Reelin: p = 0.02<br>CORT/Vehicle - CORT/Repeated Reelin: p = 0.006<br>CORT/Reelin - CORT/Repeated Reelin: p = 1 | Vehicle/Vehicle - Vehicle/Reelin: p = 0.93<br>Vehicle/Vehicle - CORT/Vehicle: p < 0.001<br>Vehicle/Vehicle - CORT/Reelin: p = 0.02<br>Vehicle/Vehicle - CORT/Repeated Reelin: p = 0.19<br>CORT/Vehicle - Vehicle/Reelin: p < 0.001<br>CORT/Vehicle - CORT/Reelin: p = 0.003<br>CORT/Vehicle - CORT/Repeated Reelin: p < 0.001<br>CORT/Reelin - CORT/Repeated Reelin: p = 0.97 | 1.1<br>3.05<br>1.09<br>0.87<br>1.95<br>1.96<br>2.18<br>0.22 | 0.63<br>4.09<br>1.87<br>1.32<br>3.46<br>2.23<br>2.77<br>0.54 | | |
| 3B | Behavioural results - Novel object location test | Three-way ANOVA<br>One-sample t tests against DI = 0 | Vehicle/Vehicle = 6<br>Vehicle/Reelin = 6<br>CORT/Vehicle = 6<br>CORT/Reelin = 6<br>CORT/Repeated Reelin = 6 | CORT: p = 0.98<br>Reelin: p = 0.95<br>Sex: p = 0.04<br>CORT:Reelin: p = 0.56<br>CORT:Sex: p = 0.27<br>Reelin:Sex: p = 0.21<br>CORT:Reelin:Sex: p = 0.51 | F(1,48) = 0.0006<br>F(2,48) = 0.05<br>F(1,48) = 4.31<br>F(1,48) = 0.34<br>F(1,48) =1.27<br>F(2,48) = 1.60<br>F(1,48) = 0.45 | 1.13053E-05<br>0.001957639<br>0.079363096<br>0.006822672<br>0.024712639<br>0.060177159<br>0.008838771 | | **t test p value**<br>VV males: p = 0.03<br>VR males: p = 0.09<br>CV males: p = 0.005<br>CR males: p = 0.15<br>CRR males: p = 0.14<br>VV females: p = 0.06<br>VR females: p = 0.54<br>CV females: p = 0.07<br>CR females: p = 0.03<br>CRR females: p = 0.07 | discrepancy<br>VV males: 0.37<br>VR males: 0.29<br>CV males: 0.35<br>CR males: 0.25<br>CRR males: 0.16<br>VV females:0.13<br>VR females: 0.08<br>CV females: 0.07<br>CR females: 0.21<br>CRR females: 0.27 | 95% CI<br>VV males: [0.04,0.70]<br>VR males: [-0.06,0.64]<br>CV males: [0.16,0.54]<br>CR males: [-0.12,0.63]<br>CRR males: [-0.08,0.41]<br>VV females:[-0.007,0.26]<br>VR females: [-0.23,0.39]<br>CV females: [-0.009,0.14]<br>CR females: [0.02,0.38]<br>CRR females: [-0.03,0.57] | | | | |
| 3Ca | Behavioural results - Open field test (Time in Center Zone) | Three-way ANOVA<br>Post hoc:<br>Šídák's multiple comparisons test | Vehicle/Vehicle = 6<br>Vehicle/Reelin = 6<br>CORT/Vehicle = 6<br>CORT/Reelin = 6<br>CORT/Repeated Reelin = 6 | CORT: p = 0.73<br>Reelin: p = 0.85<br>Sex: p = 0.69<br>CORT:Reelin: p = 0.71<br>CORT:Sex: p = 0.50<br>Reelin:Sex: p = 0.98<br>CORT:Reelin:Sex: p = 0.85 | F(1,48) = 0.12<br>F(2,48) = 0.16<br>F(1,48) = 0.16<br>F(1,48) = 0.14<br>F(1,48) = 0.47<br>F(2,48) = 0.02<br>F(1,48) = 0.04 | 0.002489373<br>0.006520023<br>0.003245029<br>0.002848985<br>0.00932228<br>0.000688344<br>0.00071535 | | | | | | | | |
| 3Cb | Behavioural results - Open field test (Frequency of Center Zone Visits) | Three-way ANOVA<br>Post hoc:<br>Šídák's multiple comparisons test | Vehicle/Vehicle = 6<br>Vehicle/Reelin = 6<br>CORT/Vehicle = 6<br>CORT/Reelin = 6<br>CORT/Repeated Reelin = 6 | CORT: p = 0.24<br>Reelin: p = 0.79<br>Sex: p = 0.60<br>CORT:Reelin: p = 0.89<br>CORT:Sex: p = 0.44<br>Reelin:Sex: p = 0.78<br>CORT:Reelin:Sex: p = 0.82 | F(1,48) = 1.43<br>F(2,48) = 0.23<br>F(1,48) = 0.28<br>F(1,48) = 0.02<br>F(1,48) = 0.60<br>F(2,48) = 0.25<br>F(1,48) = 0.05 | 0.027753753<br>0.009254272<br>0.005508026<br>0.000411678<br>0.011885202<br>0.010024463<br>0.001053221 | | | | | | | | |
| 3Cc | Behavioural results - Open field test (Latency to Enter Center) | Three-way ANOVA<br>Post hoc:<br>Šídák's multiple comparisons test | Vehicle/Vehicle = 6<br>Vehicle/Reelin = 6<br>CORT/Vehicle = 6<br>CORT/Reelin = 6<br>CORT/Repeated Reelin = 6 | CORT: p = 0.30<br>Reelin: p = 0.87<br>Sex: p = 0.90<br>CORT:Reelin: p = 0.86<br>CORT:Sex: p = 0.83<br>Reelin:Sex: p = 0.72<br>CORT:Reelin:Sex: p = 0.36 | F(1,48) = 1.08<br>F(2,48) = 0.14<br>F(1,48) = 0.02<br>F(1,48) = 0.03<br>F(1,48) = 0.05<br>F(2,48) = 0.33<br>F(1,48) = 0.84 | 0.022404651<br>0.005766806<br>0.000337868<br>0.000644464<br>0.000986959<br>0.013665326<br>0.017520421 | | | | | | | | |
| 3Cd | Behavioural results - Open field test (Velocity) | Three-way ANOVA<br>Post hoc:<br>Šídák's multiple comparisons test | Vehicle/Vehicle = 6<br>Vehicle/Reelin = 6<br>CORT/Vehicle = 6<br>CORT/Reelin = 6<br>CORT/Repeated Reelin = 6 | CORT: p = 0.74<br>Reelin: p = 0.33<br>Sex: p = 0.03<br>CORT:Reelin: p = 0.25<br>CORT:Sex: p = 0.58<br>Reelin:Sex: p = 0.51<br>CORT:Reelin:Sex: p = 0.29 | F(1,48) = 0.11<br>F(2,48) = 1.14<br>F(1,48) = 4.81<br>F(1,48) = 1.33<br>F(1,48) = 0.30<br>F(2,48) = 0.69<br>F(1,48) = 1.14 | 0.002201752<br>0.043753309<br>0.087850123<br>0.025923261<br>0.006017652<br>0.02686523<br>0.022380189 | | | Vehicle/Vehicle - Vehicle/Reelin: p = 0.80<br>Vehicle/Vehicle - CORT/Vehicle: p = 1<br>Vehicle/Vehicle - CORT/Reelin: p = 1<br>Vehicle/Vehicle - CORT/Repeated Reelin: p = 0.94<br>CORT/Vehicle - Vehicle/Reelin: p = 0.95<br>CORT/Vehicle - CORT/Reelin: p = 0.98<br>CORT/Vehicle - CORT/Repeated Reelin: p = 0.99<br>CORT/Reelin - CORT/Repeated Reelin: p = 0.63 | Vehicle/Vehicle - Vehicle/Reelin: p = 0.99<br>Vehicle/Vehicle - CORT/Vehicle: p = 1<br>Vehicle/Vehicle - CORT/Reelin: p = 0.97<br>Vehicle/Vehicle - CORT/Repeated Reelin: p = 1<br>CORT/Vehicle - Vehicle/Reelin: p = 0.99<br>CORT/Vehicle - CORT/Reelin: p = 0.98<br>CORT/Vehicle - CORT/Repeated Reelin: p = 1<br>CORT/Reelin - CORT/Repeated Reelin: p = 0.96 | 0.78<br>0.19<br>0.32<br>0.6<br>0.6<br>0.5<br>0.41<br>0.92 | 0.45<br>0.02<br>0.53<br>0.04<br>0.43<br>0.5<br>0.07<br>0.57 | | |
| 3Ce | Behavioural results - Open field test (Distance Travelled) | Three-way ANOVA<br>Post hoc:<br>Šídák's multiple comparisons test | Vehicle/Vehicle = 6<br>Vehicle/Reelin = 6<br>CORT/Vehicle = 6<br>CORT/Reelin = 6<br>CORT/Repeated Reelin = 6 | CORT: p = 0.74<br>Reelin: p = 0.33<br>Sex: p = 0.03<br>CORT:Reelin: p = 0.25<br>CORT:Sex: p = 0.58<br>Reelin:Sex: p = 0.50<br>CORT:Reelin:Sex: p = 0.29 | F(1,48) = 0.11<br>F(2,48) = 1.14<br>F(1,48) = 4.84<br>F(1,48) = 1.34<br>F(1,48) = 0.31<br>F(2,48) = 0.70<br>F(1,48) = 1.15 | 0.002207376<br>0.043740589<br>0.088323576<br>0.026131889<br>0.006083321<br>0.027269857<br>0.022419553 | | | Vehicle/Vehicle - Vehicle/Reelin: p = 0.79<br>Vehicle/Vehicle - CORT/Vehicle: p = 1<br>Vehicle/Vehicle - CORT/Reelin: p = 1<br>Vehicle/Vehicle - CORT/Repeated Reelin: p = 0.94<br>CORT/Vehicle - Vehicle/Reelin: p = 0.95<br>CORT/Vehicle - CORT/Reelin: p = 0.98<br>CORT/Vehicle - CORT/Repeated Reelin: p = 0.99<br>CORT/Reelin - CORT/Repeated Reelin: p = 0.63 | Vehicle/Vehicle - Vehicle/Reelin: p = 0.99<br>Vehicle/Vehicle - CORT/Vehicle: p = 1<br>Vehicle/Vehicle - CORT/Reelin: p = 0.97<br>Vehicle/Vehicle - CORT/Repeated Reelin: p = 1<br>CORT/Vehicle - Vehicle/Reelin: p = 0.99<br>CORT/Vehicle - CORT/Reelin: p = 0.98<br>CORT/Vehicle - CORT/Repeated Reelin: p = 1<br>CORT/Reelin - CORT/Repeated Reelin: p = 0.96 | 0.79<br>0.19<br>0.31<br>0.6<br>0.6<br>0.5<br>0.42<br>0.92 | 0.45<br>0.02<br>0.53<br>0.04<br>0.43<br>0.5<br>0.07<br>0.57 | | |

| Figure Number | Figure Description | Statistical Test | n | p value | F values | Partial Eta Squared | R Squared | Post-Hocs | Post-Hocs Females | Post-Hocs Males | Cohen's d Fem | Cohen's d Mal | Westfall d Females | Westfall d Males |
|---|---|---|---|---|---|---|---|---|---|---|---|---|---|---|
| 4A | Microglia density - Polymorphic layer | Three-way ANOVA<br>Post hoc: Šídák's multiple comparisons test | Vehicle/Vehicle = 6<br>Vehicle/Reelin = 6<br>CORT/Vehicle = 6<br>CORT/Reelin = 6<br>CORT/Repeated Reelin = 6 | CORT: p = 0.01<br>Reelin: p = 0.48<br>Sex: p = 0.93<br>CORT:Reelin: p = 0.19<br>CORT:Sex: p = 0.37<br>Reelin:Sex: p = 0.57<br>CORT:Reelin:Sex: p = 0.12 | F(1,48) = 6.60<br>F(2,48) = 0.74<br>F(1,48) = 0.008<br>F(1,48) = 1.74<br>F(1,48) = 0.83<br>F(2,48) = 0.58<br>F(1,48) = 2.46 | 0.11665118<br>0.028681378<br>0.000165017<br>0.033706951<br>0.016411508<br>0.022537244<br>0.046972797 | | | Vehicle/Vehicle - Vehicle/Reelin: p = 0.95<br>Vehicle/Vehicle - CORT/Vehicle: p = 0.99<br>Vehicle/Vehicle - CORT/Reelin: p = 1<br>Vehicle/Vehicle - CORT/Repeated Reelin: p = 1<br>CORT/Vehicle - Vehicle/Reelin: p = 0.46<br>CORT/Vehicle - CORT/Reelin: p = 0.86<br>CORT/Vehicle - CORT/Repeated Reelin: p = 1<br>CORT/Reelin - CORT/Repeated Reelin: p = 0.99 | Vehicle/Vehicle - Vehicle/Reelin: p = 0.70<br>Vehicle/Vehicle - CORT/Vehicle: p = 1<br>Vehicle/Vehicle - CORT/Reelin: p = 0.71<br>Vehicle/Vehicle - CORT/Repeated Reelin: p = 0.97<br>CORT/Vehicle - Vehicle/Reelin: p = 0.63<br>CORT/Vehicle - CORT/Reelin: p = 0.78<br>CORT/Vehicle - CORT/Repeated Reelin: p = 0.98<br>CORT/Reelin - CORT/Repeated Reelin: p = 1 | 0.58<br>0.47<br>0.25<br>0.22<br>1.05<br>0.72<br>0.25<br>0.47 | 0.87<br>0.05<br>0.86<br>0.54<br>0.92<br>0.8<br>0.49<br>0.31 | | |
| 4B | Microglia density - Subgranular zone | Three-way ANOVA<br>Post hoc: Šídák's multiple comparisons test | Vehicle/Vehicle = 6<br>Vehicle/Reelin = 6<br>CORT/Vehicle = 6<br>CORT/Reelin = 6<br>CORT/Repeated Reelin = 6 | CORT: p = 0.65<br>Reelin: p = 0.84<br>Sex: p = 0.50<br>CORT:Reelin: p = 0.47<br>CORT:Sex: p = 0.22<br>Reelin:Sex: p = 0.91<br>CORT:Reelin:Sex: p = 0.49 | F(1,48) = 0.20<br>F(2,48) = 0.18<br>F(1,48) = 0.46<br>F(1,48) = 0.53<br>F(1,48) = 1.55<br>F(2,48) = 0.09<br>F(1,48) = 0.48 | 0.004039985<br>0.007090397<br>0.009168711<br>0.010527938<br>0.030049945<br>0.003690485<br>0.009581111 | | | | | | | | |

| Figure Number | Figure Description | Statistical Test | n | p value | F values | R Squared | Post-Hocs Females | Post-Hocs Males |
|---|---|---|---|---|---|---|---|---|
| 5Aa | Microglia process tracings - Length 0-10 um) | Nested three-way ANOVA<br><br>Post hoc: Šídák's multiple comparisons test | Vehicle/Vehicle = 6<br>Vehicle/Reelin = 6<br>CORT/Vehicle = 6<br>CORT/Reelin = 6<br>CORT/Repeated | CORT: p = 0.08<br>Reelin: p = 1<br>Sex: p = 0.34<br>CORT:Reelin: p = 0.31<br>CORT:Sex: p = 0.16<br>Reelin:Sex: p = 0.23<br>CORT:Reelin:Sex: p = 0.12 | F(1,48) = 3.16<br>F(2,48) = 0.004<br>F(1,48) = 0.92<br>F(1,48) = 1.06<br>F(1,48) = 2.03<br>F(2,48) = 1.53<br>F(1,48) = 2.44 | Marginal R^2 = 0.06<br>Conditional R^2 = 0.21 | | |
| 5Ab | Microglia process tracings - Length 10-20 um) | Nested three-way ANOVA<br><br>Post hoc: Šídák's multiple comparisons test | Vehicle/Vehicle = 6<br>Vehicle/Reelin = 6<br>CORT/Vehicle = 6<br>CORT/Reelin = 6<br>CORT/Repeated | CORT: p = 0.13<br>Reelin: p = 0.98<br>Sex: p = 0.26<br>CORT:Reelin: p = 0.12<br>CORT:Sex: p = 0.69<br>Reelin:Sex: p = 0.39<br>CORT:Reelin:Sex: p = 1 | F(1,48) = 2.32<br>F(2,48) = 0.02<br>F(1,48) = 1.32<br>F(1,48) = 2.47<br>F(1,48) = 0.17<br>F(2,48) = 0.96<br>F(1,48) < 0.001 | Marginal R^2 = 0.04<br>Conditional R^2 = 0.20 | | |
| 5Ac | Microglia process tracings - Length 20-30 um) | Nested three-way ANOVA<br><br>Post hoc: Šídák's multiple comparisons test | Vehicle/Vehicle = 6<br>Vehicle/Reelin = 6<br>CORT/Vehicle = 6<br>CORT/Reelin = 6<br>CORT/Repeated | CORT: p = 0.07<br>Reelin: p = 0.85<br>Sex: p = 0.06<br>CORT:Reelin: p = 0.15<br>CORT:Sex: p = 0.70<br>Reelin:Sex: p = 0.27<br>CORT:Reelin:Sex: p = 0.96 | F(1,48) = 3.52<br>F(2,48) = 0.16<br>F(1,48) = 3.71<br>F(1,48) = 2.19<br>F(1,48) = 0.15<br>F(2,48) = 1.35<br>F(1,48) = 0.002 | Marginal R^2 = 0.06<br>Conditional R^2 = 0.25 | | |
| 5Ad | Microglia process tracings - Length 30-40 um) | Nested three-way ANOVA<br><br>Post hoc: Šídák's multiple comparisons test | Vehicle/Vehicle = 6<br>Vehicle/Reelin = 6<br>CORT/Vehicle = 6<br>CORT/Reelin = 6<br>CORT/Repeated | CORT: p = 0.12<br>Reelin: p = 0.36<br>Sex: p = 0.04<br>CORT:Reelin: p = 0.17<br>CORT:Sex: p = 0.45<br>Reelin:Sex: p = 0.46<br>CORT:Reelin:Sex: p = 0.86 | F(1,48) = 2.47<br>F(2,48) = 1.03<br>F(1,48) = 4.29<br>F(1,48) = 1.91<br>F(1,48) = 0.58<br>F(2,48) = 0.79<br>F(1,48) = 0.03 | Marginal R^2 = 0.06<br>Conditional R^2 = 0.26 | Vehicle/Vehicle - Vehicle/Reelin: p = 0.24<br>Vehicle/Vehicle - CORT/Vehicle: p = 0.07<br>Vehicle/Vehicle - CORT/Reelin: p = 0.10<br>Vehicle/Vehicle - CORT/Repeated Reelin: p = 0.02<br>CORT/Vehicle - Vehicle/Reelin: p = 0.17<br>CORT/Vehicle - CORT/Reelin: p = 0.17<br>CORT/Vehicle - CORT/Repeated Reelin: p = 0.05<br>CORT/Reelin - CORT/Repeated Reelin: p = 0.12 | Vehicle/Vehicle - Vehicle/Reelin: p = 0.31<br>Vehicle/Vehicle - CORT/Vehicle: p = 0.13<br>Vehicle/Vehicle - CORT/Reelin: p = 0.36<br>Vehicle/Vehicle - CORT/Repeated Reelin: p = 0.31<br>CORT/Vehicle - Vehicle/Reelin: p = 0.44<br>CORT/Vehicle - CORT/Reelin: p = 0.22<br>CORT/Vehicle - CORT/Repeated Reelin: p = 0.44<br>CORT/Reelin - CORT/Repeated Reelin: p = 0.67 |
| 5Ae | Microglia process tracings - Length 40-50 um) | Nested three-way ANOVA<br><br>Post hoc: Šídák's multiple comparisons test | Vehicle/Vehicle = 6<br>Vehicle/Reelin = 6<br>CORT/Vehicle = 6<br>CORT/Reelin = 6<br>CORT/Repeated | CORT: p = 0.16<br>Reelin: p = 0.45<br>Sex: p = 0.14<br>CORT:Reelin: p = 0.85<br>CORT:Sex: p = 0.69<br>Reelin:Sex: p = 0.88<br>CORT:Reelin:Sex: p = 0.58 | F(1,48) = 1.99<br>F(2,48) = 0.82<br>F(1,48) = 2.28<br>F(1,48) = 0.04<br>F(1,48) = 0.16<br>F(2,48) = 0.13<br>F(1,48) = 0.32 | Marginal R^2 = 0.03<br>Conditional R^2 = 0.17 | | |
| 5Af | Microglia process tracings - Total Length | Nested three-way ANOVA<br><br>Post hoc: Šídák's multiple comparisons test | Vehicle/Vehicle = 6<br>Vehicle/Reelin = 6<br>CORT/Vehicle = 6<br>CORT/Reelin = 6<br>CORT/Repeated | CORT: p = 0.05<br>Reelin: p = 0.85<br>Sex: p = 0.06<br>CORT:Reelin: p = 0.11<br>CORT:Sex: p = 0.48<br>Reelin:Sex: p = 0.28<br>CORT:Reelin:Sex: p = 0.90 | F(1,48) = 4.16<br>F(2,48) = 0.17<br>F(1,48) = 3.83<br>F(1,48) = 2.62<br>F(1,48) = 0.51<br>F(2,48) = 1.29<br>F(1,48) = 0.02 | Marginal R^2 = 0.07<br>Conditional R^2 = 0.29 | Vehicle/Vehicle - Vehicle/Reelin: p = 0.22<br>Vehicle/Vehicle - CORT/Vehicle: p = 0.07<br>Vehicle/Vehicle - CORT/Reelin: p = 0.32<br>Vehicle/Vehicle - CORT/Repeated Reelin: p = 0.31<br>CORT/Vehicle - Vehicle/Reelin: p = 0.14<br>CORT/Vehicle - CORT/Reelin: p = 0.39<br>CORT/Vehicle - CORT/Repeated Reelin: p = 0.38<br>CORT/Reelin - CORT/Repeated Reelin: p = 0.006 | Vehicle/Vehicle - Vehicle/Reelin: p = 0.44<br>Vehicle/Vehicle - CORT/Vehicle: p = 0.22<br>Vehicle/Vehicle - CORT/Reelin: p = 0.30<br>Vehicle/Vehicle - CORT/Repeated Reelin: p = 0.15<br>CORT/Vehicle - Vehicle/Reelin: p = 0.66<br>CORT/Vehicle - CORT/Reelin: p = 0.08<br>CORT/Vehicle - CORT/Repeated Reelin: p = 0.37<br>CORT/Reelin - CORT/Repeated Reelin: p = 0.45 |

| Figure Number | Figure Description | Statistical Test | n | p value | F values | R Squared | Post-Hocs Females | Post-Hocs Males |
|---|---|---|---|---|---|---|---|---|
| 5Ba | Microglia process tracings - Endings 0-10 um) | Nested three-way ANOVA<br><br>Post hoc: Šídák's multiple comparisons test | Vehicle/Vehicle = 6<br>Vehicle/Reelin = 6<br>CORT/Vehicle = 6<br>CORT/Reelin = 6<br>CORT/Repeated | CORT: p = 0.82<br>Reelin: p = 0.37<br>Sex: p = 0.16<br>CORT:Reelin: p = 0.82<br>CORT:Sex: p = 0.17<br>Reelin:Sex: p = 0.07<br>CORT:Reelin:Sex: p = 0.39 | F(1,48) = 0.05<br>F(2,48) = 1.02<br>F(1,48) = 2.002<br>F(1,48) = 0.05<br>F(1,48) = 1.91<br>F(2,48) = 2.79<br>F(1,48) = 0.75 | Marginal $R^2$ = 0.05<br>Conditional $R^2$ = 0.19 | | |
| 5Bb | Microglia process tracings - Endings 10-20 um) | Nested three-way ANOVA<br><br>Post hoc: Šídák's multiple comparisons test | Vehicle/Vehicle = 6<br>Vehicle/Reelin = 6<br>CORT/Vehicle = 6<br>CORT/Reelin = 6<br>CORT/Repeated | CORT: p = 0.17<br>Reelin: p = 0.65<br>Sex: p = 0.47<br>CORT:Reelin: p = 0.77<br>CORT:Sex: p = 0.29<br>Reelin:Sex: p = 0.90<br>CORT:Reelin:Sex: p = 0.59 | F(1,48) = 1.90<br>F(2,48) = 0.43<br>F(1,48) = 0.54<br>F(1,48) = 0.09<br>F(1,48) = 1.15<br>F(2,48) = 0.11<br>F(1,48) = 0.29 | Marginal $R^2$ = 0.03<br>Conditional $R^2$ = 0.17 | | |
| 5Bc | Microglia process tracings - Endings 20-30 um) | Nested three-way ANOVA<br><br>Post hoc: Šídák's multiple comparisons test | Vehicle/Vehicle = 6<br>Vehicle/Reelin = 6<br>CORT/Vehicle = 6<br>CORT/Reelin = 6<br>CORT/Repeated | CORT: p = 0.08<br>Reelin: p = 0.71<br>Sex: p = 0.04<br>CORT:Reelin: p = 0.17<br>CORT:Sex: p = 0.39<br>Reelin:Sex: p = 0.28<br>CORT:Reelin:Sex: p = 0.28 | F(1,48) = 3.27<br>F(2,48) = 0.34<br>F(1,48) = 4.33<br>F(1,48) = 1.90<br>F(1,48) = 0.76<br>F(2,48) = 1.30<br>F(1,48) = 1.19 | Marginal $R^2$ = 0.07<br>Conditional $R^2$ = 0.22 | Vehicle/Vehicle - Vehicle/Reelin: p = 0.21<br>Vehicle/Vehicle - CORT/Vehicle: p = 0.24<br>Vehicle/Vehicle - CORT/Reelin: p = 0.33<br>Vehicle/Vehicle - CORT/Repeated Reelin: p = 0.33<br>CORT/Vehicle - Vehicle/Reelin: p = 0.03<br>CORT/Vehicle - CORT/Reelin: p = 0.57<br>CORT/Vehicle - CORT/Repeated Reelin: p = 0.57<br>CORT/Reelin - CORT/Repeated Reelin: p < 0.001 | Vehicle/Vehicle - Vehicle/Reelin: p = 0.33<br>Vehicle/Vehicle - CORT/Vehicle: p = 0.38<br>Vehicle/Vehicle - CORT/Reelin: p = 0.14<br>Vehicle/Vehicle - CORT/Repeated Reelin: p = 0.27<br>CORT/Vehicle - Vehicle/Reelin: p = 0.71<br>CORT/Vehicle - CORT/Reelin: p = 0.24<br>CORT/Vehicle - CORT/Repeated Reelin: p = 0.11<br>CORT/Reelin - CORT/Repeated Reelin: p = 0.13 |
| 5Bd | Microglia process tracings - Endings 30-40 um) | Nested three-way ANOVA<br><br>Post hoc: Šídák's multiple comparisons test | Vehicle/Vehicle = 6<br>Vehicle/Reelin = 6<br>CORT/Vehicle = 6<br>CORT/Reelin = 6<br>CORT/Repeated | CORT: p = 0.09<br>Reelin: p = 0.35<br>Sex: p = 0.10<br>CORT:Reelin: p = 0.14<br>CORT:Sex: p = 0.77<br>Reelin:Sex: p = 0.27<br>CORT:Reelin:Sex: p = 0.80 | F(1,48) = 2.91<br>F(2,48) = 1.09<br>F(1,48) = 2.80<br>F(1,48) = 2.19<br>F(1,48) = 0.08<br>F(2,48) = 1.34<br>F(1,48) = 0.07 | Marginal $R^2$ = 0.05<br>Conditional $R^2$ = 0.17 | | |
| 5Be | Microglia process tracings - Endings 40-50 um) | Nested three-way ANOVA<br><br>Post hoc: Šídák's multiple comparisons test | Vehicle/Vehicle = 6<br>Vehicle/Reelin = 6<br>CORT/Vehicle = 6<br>CORT/Reelin = 6<br>CORT/Repeated | CORT: p = 0.26<br>Reelin: p = 0.49<br>Sex: p = 0.10<br>CORT:Reelin: p = 0.60<br>CORT:Sex: p = 0.84<br>Reelin:Sex: p = 0.37<br>CORT:Reelin:Sex: p = 0.60 | F(1,48) = 1.29<br>F(2,48) = 0.73<br>F(1,48) = 2.87<br>F(1,48) = 0.29<br>F(1,48) = 0.04<br>F(2,48) = 1.02<br>F(1,48) = 0.29 | Marginal $R^2$ = 0.03<br>Conditional $R^2$ = 0.18 | | |
| 5Bf | Microglia process tracings - Total Endings | Nested three-way ANOVA<br><br>Post hoc: Šídák's multiple comparisons test | Vehicle/Vehicle = 6<br>Vehicle/Reelin = 6<br>CORT/Vehicle = 6<br>CORT/Reelin = 6<br>CORT/Repeated | CORT: p = 0.06<br>Reelin: p = 0.95<br>Sex: p = 0.04<br>CORT:Reelin: p = 0.30<br>CORT:Sex: p = 0.29<br>Reelin:Sex: p = 0.31<br>CORT:Reelin:Sex: p = 0.54 | F(1,48) = 3.63<br>F(2,48) = 0.06<br>F(1,48) = 4.48<br>F(1,48) = 1.08<br>F(1,48) = 1.17<br>F(2,48) = 1.20<br>F(1,48) = 0.39 | Marginal $R^2$ = 0.07<br>Conditional $R^2$ = 0.30 | Vehicle/Vehicle - Vehicle/Reelin: p = 0.14<br>Vehicle/Vehicle - CORT/Vehicle: p = 0.15<br>Vehicle/Vehicle - CORT/Reelin: p = 0.29<br>Vehicle/Vehicle - CORT/Repeated Reelin: p = 0.31<br>CORT/Vehicle - Vehicle/Reelin: p = 0.003<br>CORT/Vehicle - CORT/Reelin: p = 0.44<br>CORT/Vehicle - CORT/Repeated Reelin: p = 0.46<br>CORT/Reelin - CORT/Repeated Reelin: p = 0.017 | Vehicle/Vehicle - Vehicle/Reelin: p = 0.33<br>Vehicle/Vehicle - CORT/Vehicle: p = 0.45<br>Vehicle/Vehicle - CORT/Reelin: p = 0.26<br>Vehicle/Vehicle - CORT/Repeated Reelin: p = 0.16<br>CORT/Vehicle - Vehicle/Reelin: p = 0.78<br>CORT/Vehicle - CORT/Reelin: p = 0.19<br>CORT/Vehicle - CORT/Repeated Reelin: p = 0.29<br>CORT/Reelin - CORT/Repeated Reelin: p = 0.10 |

| Figure Number | Figure Description | Statistical Test | n | p value | F values | R Squared | Post-Hocs Females | Post-Hocs Males |
|---|---|---|---|---|---|---|---|---|
| 5Ca | Microglia process tracings - Intersections 0-10 um) | Nested three-way ANOVA<br><br>Post hoc: Šídák's multiple comparisons test | Vehicle/Vehicle = 6<br>Vehicle/Reelin = 6<br>CORT/Vehicle = 6<br>CORT/Reelin = 6<br>CORT/Repeated | CORT: p = 0.007<br>Reelin: p = 0.84<br>Sex: p = 0.43<br>CORT:Reelin: p = 0.61<br>CORT:Sex: p = 0.50<br>Reelin:Sex: p = 0.73<br>CORT:Reelin:Sex: p = 0.56 | F(1,48) = 7.85<br>F(2,48) = 0.18<br>F(1,48) = 0.63<br>F(1,48) = 0.26<br>F(1,48) = 0.47<br>F(2,48) = 0.32<br>F(1,48) = 0.34 | Marginal R^2 = 0.04<br>Conditional R^2 = 0.16 | Vehicle/Vehicle - Vehicle/Reelin: p = 0.03<br>Vehicle/Vehicle - CORT/Vehicle: p = 0.15<br>Vehicle/Vehicle - CORT/Reelin: p = 0.45<br>Vehicle/Vehicle - CORT/Repeated Reelin: p = 0.23<br>CORT/Vehicle - Vehicle/Reelin: p = 0.18<br>CORT/Vehicle - CORT/Reelin: p = 0.30<br>CORT/Vehicle - CORT/Repeated Reelin: p = 0.07<br>CORT/Reelin - CORT/Repeated Reelin: p = 0.23 | Vehicle/Vehicle - Vehicle/Reelin: p = 0.09<br>Vehicle/Vehicle - CORT/Vehicle: p = 0.53<br>Vehicle/Vehicle - CORT/Reelin: p = 0.42<br>Vehicle/Vehicle - CORT/Repeated Reelin: p = 0.34<br>CORT/Vehicle - Vehicle/Reelin: p = 0.63<br>CORT/Vehicle - CORT/Reelin: p = 0.11<br>CORT/Vehicle - CORT/Repeated Reelin: p = 0.19<br>CORT/Reelin - CORT/Repeated Reelin: p = 0.08 |
| 5Cb | Microglia process tracings - Intersections 10-20 um) | Nested three-way ANOVA<br><br>Post hoc: Šídák's multiple comparisons test | Vehicle/Vehicle = 6<br>Vehicle/Reelin = 6<br>CORT/Vehicle = 6<br>CORT/Reelin = 6<br>CORT/Repeated | CORT: p = 0.18<br>Reelin: p = 1<br>Sex: p = 0.17<br>CORT:Reelin: p = 0.07<br>CORT:Sex: p = 0.68<br>Reelin:Sex: p = 0.23<br>CORT:Reelin:Sex: p = 0.98 | F(1,48) = 1.88<br>F(2,48) = 0.0007<br>F(1,48) = 1.90<br>F(1,48) = 3.47<br>F(1,48) = 0.17<br>F(2,48) = 1.52<br>F(1,48) = 0.0006 | Marginal R^2 = 0.05<br>Conditional R^2 = 0.19 | | |
| 5Cc | Microglia process tracings - Intersections 20-30 um) | Nested three-way ANOVA<br><br>Post hoc: Šídák's multiple comparisons test | Vehicle/Vehicle = 6<br>Vehicle/Reelin = 6<br>CORT/Vehicle = 6<br>CORT/Reelin = 6<br>CORT/Repeated | CORT: p = 0.03<br>Reelin: p = 0.18<br>Sex: p = 0.20<br>CORT:Reelin: p = 0.23<br>CORT:Sex: p = 0.99<br>Reelin:Sex: p = 0.23<br>CORT:Reelin:Sex: p = 0.70 | F(1,48) = 4.73<br>F(2,48) = 1.78<br>F(1,48) = 1.65<br>F(1,48) = 1.47<br>F(1,48) = 0.0003<br>F(2,48) = 1.52<br>F(1,48) = 0.15 | Marginal R^2 = 0.06<br>Conditional R^2 = 0.19 | Vehicle/Vehicle - Vehicle/Reelin: p = 0.17<br>Vehicle/Vehicle - CORT/Vehicle: p = 0.21<br>Vehicle/Vehicle - CORT/Reelin: p = 0.29<br>Vehicle/Vehicle - CORT/Repeated Reelin: p = 0.24<br>CORT/Vehicle - Vehicle/Reelin: p = 0.38<br>CORT/Vehicle - CORT/Reelin: p = 0.08<br>CORT/Vehicle - CORT/Repeated Reelin: p = 0.03<br>CORT/Reelin - CORT/Repeated Reelin: p = 0.05 | Vehicle/Vehicle - Vehicle/Reelin: p = 0.42<br>Vehicle/Vehicle - CORT/Vehicle: p = 0.09<br>Vehicle/Vehicle - CORT/Reelin: p = 0.16<br>Vehicle/Vehicle - CORT/Repeated Reelin: p = 0.52<br>CORT/Vehicle - Vehicle/Reelin: p = 0.51<br>CORT/Vehicle - CORT/Reelin: p = 0.07<br>CORT/Vehicle - CORT/Repeated Reelin: p = 0.62<br>CORT/Reelin - CORT/Repeated Reelin: p = 0.69 |
| 5Cd | Microglia process tracings - Intersections 30-40 um) | Nested three-way ANOVA<br><br>Post hoc: Šídák's multiple comparisons test | Vehicle/Vehicle = 6<br>Vehicle/Reelin = 6<br>CORT/Vehicle = 6<br>CORT/Reelin = 6<br>CORT/Repeated | CORT: p = 0.16<br>Reelin: p = 0.34<br>Sex: p = 0.11<br>CORT:Reelin: p = 0.45<br>CORT:Sex: p = 0.78<br>Reelin:Sex: p = 0.59<br>CORT:Reelin:Sex: p = 0.98 | F(1,48) = 2.07<br>F(2,48) = 1.09<br>F(1,48) = 2.69<br>F(1,48) = 0.59<br>F(1,48) = 0.08<br>F(2,48) = 0.53<br>F(1,48) = 0.0009 | Marginal R^2 = 0.03<br>Conditional R^2 = 0.18 | | |
| 5Ce | Microglia process tracings - Total Intersections | Nested three-way ANOVA<br><br>Post hoc: Šídák's multiple comparisons test | Vehicle/Vehicle = 6<br>Vehicle/Reelin = 6<br>CORT/Vehicle = 6<br>CORT/Reelin = 6<br>CORT/Repeated | CORT: p = 0.02<br>Reelin: p = 0.60<br>Sex: p = 0.14<br>CORT:Reelin: p = 0.15<br>CORT:Sex: p = 0.66<br>Reelin:Sex: p = 0.33<br>CORT:Reelin:Sex: p = 0.96 | F(1,48) = 5.60<br>F(2,48) = 0.51<br>F(1,48) = 2.28<br>F(1,48) = 2.13<br>F(1,48) = 0.19<br>F(2,48) = 1.15<br>F(1,48) = 0.003 | Marginal R^2 = 0.06<br>Conditional R^2 = 0.26 | Vehicle/Vehicle - Vehicle/Reelin: p = 0.15<br>Vehicle/Vehicle - CORT/Vehicle: p = 0.07<br>Vehicle/Vehicle - CORT/Reelin: p = 0.42<br>Vehicle/Vehicle - CORT/Repeated Reelin: p = 0.31<br>CORT/Vehicle - Vehicle/Reelin: p = 0.22<br>CORT/Vehicle - CORT/Reelin: p = 0.35<br>CORT/Vehicle - CORT/Repeated Reelin: p = 0.24<br>CORT/Reelin - CORT/Repeated Reelin: p = 0.12 | Vehicle/Vehicle - Vehicle/Reelin: p = 0.40<br>Vehicle/Vehicle - CORT/Vehicle: p = 0.24<br>Vehicle/Vehicle - CORT/Reelin: p = 0.31<br>Vehicle/Vehicle - CORT/Repeated Reelin: p = 0.20<br>CORT/Vehicle - Vehicle/Reelin: p = 0.64<br>CORT/Vehicle - CORT/Reelin: p = 0.07<br>CORT/Vehicle - CORT/Repeated Reelin: p = 0.44<br>CORT/Reelin - CORT/Repeated Reelin: p = 0.51 |

| Figure Number | Figure Description | Statistical Test | n | p value | F values | R Squared | Post-Hocs Females | Post-Hocs Males |
|---|---|---|---|---|---|---|---|---|
| 5Da | Microglia process tracings - Nodes 0-10 um) | Nested three-way ANOVA<br><br>Post hoc: Šídák's multiple comparisons test | Vehicle/Vehicle = 6<br>Vehicle/Reelin = 6<br>CORT/Vehicle = 6<br>CORT/Reelin = 6<br>CORT/Repeated | CORT: p = 0.07<br>Reelin: p = 0.85<br>Sex: p = 0.03<br>CORT:Reelin: p = 0.85<br>CORT:Sex: p = 0.47<br>Reelin:Sex: p = 0.62<br>CORT:Reelin:Sex: p = 0.29 | F(1,48) = 3.43<br>F(2,48) = 0.17<br>F(1,48) = 4.86<br>F(1,48) = 0.04<br>F(1,48) = 0.54<br>F(2,48) = 0.48<br>F(1,48) = 1.17 | Marginal R^2 = 0.05<br>Conditional R^2 = 0.20 | Vehicle/Vehicle - Vehicle/Reelin: p = 0.08<br>Vehicle/Vehicle - CORT/Vehicle: p = 0.04<br>Vehicle/Vehicle - CORT/Reelin: p = 0.23<br>Vehicle/Vehicle - CORT/Repeated Reelin: p = 0.19<br>CORT/Vehicle - Vehicle/Reelin: p = 0.12<br>CORT/Vehicle - CORT/Reelin: p = 0.19<br>CORT/Vehicle - CORT/Repeated Reelin: p = 0.16<br>CORT/Reelin - CORT/Repeated Reelin: p = 0.04 | Vehicle/Vehicle - Vehicle/Reelin: p = 0.04<br>Vehicle/Vehicle - CORT/Vehicle: p = 0.60<br>Vehicle/Vehicle - CORT/Reelin: p = 0.17<br>Vehicle/Vehicle - CORT/Repeated Reelin: p = 0.33<br>CORT/Vehicle - Vehicle/Reelin: p = 0.64<br>CORT/Vehicle - CORT/Reelin: p = 0.43<br>CORT/Vehicle - CORT/Repeated Reelin: p = 0.28<br>CORT/Reelin - CORT/Repeated Reelin: p = 0.16 |
| 5Db | Microglia process tracings - Nodes 10-20 um) | Nested three-way ANOVA<br><br>Post hoc: Šídák's multiple comparisons test | Vehicle/Vehicle = 6<br>Vehicle/Reelin = 6<br>CORT/Vehicle = 6<br>CORT/Reelin = 6<br>CORT/Repeated | CORT: p = 0.27<br>Reelin: p = 0.80<br>Sex: p = 0.55<br>CORT:Reelin: p = 0.38<br>CORT:Sex: p = 0.54<br>Reelin:Sex: p = 0.33<br>CORT:Reelin:Sex: p = 0.79 | F(1,48) = 1.25<br>F(2,48) = 0.23<br>F(1,48) = 0.35<br>F(1,48) = 0.80<br>F(1,48) = 0.38<br>F(2,48) = 1.14<br>F(1,48) = 0.07 | Marginal R^2 = 0.02<br>Conditional R^2 = 0.17 | | |
| 5Dc | Microglia process tracings - Nodes 20-30 um) | Nested three-way ANOVA<br><br>Post hoc: Šídák's multiple comparisons test | Vehicle/Vehicle = 6<br>Vehicle/Reelin = 6<br>CORT/Vehicle = 6<br>CORT/Reelin = 6<br>CORT/Repeated | CORT: p = 0.02<br>Reelin: p = 0.38<br>Sex: p = 0.008<br>CORT:Reelin: p = 0.28<br>CORT:Sex: p = 0.34<br>Reelin:Sex: p = 0.44<br>CORT:Reelin:Sex: p = 0.53 | F(1,48) = 5.80<br>F(2,48) = 0.99<br>F(1,48) = 7.67<br>F(1,48) = 1.19<br>F(1,48) = 0.92<br>F(2,48) = 0.83<br>F(1,48) = 0.39 | Marginal R^2 = 0.08<br>Conditional R^2 = 0.18 | Vehicle/Vehicle - Vehicle/Reelin: p = 0.31<br>Vehicle/Vehicle - CORT/Vehicle: p = 0.04<br>Vehicle/Vehicle - CORT/Reelin: p = 0.16<br>Vehicle/Vehicle - CORT/Repeated Reelin: p = 0.25<br>CORT/Vehicle - Vehicle/Reelin: p = 0.27<br>CORT/Vehicle - CORT/Reelin: p = 0.19<br>CORT/Vehicle - CORT/Repeated Reelin: p = 0.29<br>CORT/Reelin - CORT/Repeated Reelin: p = 0.10 | Vehicle/Vehicle - Vehicle/Reelin: p = 0.42<br>Vehicle/Vehicle - CORT/Vehicle: p = 0.43<br>Vehicle/Vehicle - CORT/Reelin: p = 0.15<br>Vehicle/Vehicle - CORT/Repeated Reelin: p = 0.18<br>CORT/Vehicle - Vehicle/Reelin: p = 0.84<br>CORT/Vehicle - CORT/Reelin: p = 0.28<br>CORT/Vehicle - CORT/Repeated Reelin: p = 0.24<br>CORT/Reelin - CORT/Repeated Reelin: p = 0.04 |
| 5Dd | Microglia process tracings - Nodes 30-40 um) | Nested three-way ANOVA<br><br>Post hoc: Šídák's multiple comparisons test | Vehicle/Vehicle = 6<br>Vehicle/Reelin = 6<br>CORT/Vehicle = 6<br>CORT/Reelin = 6<br>CORT/Repeated | CORT: p = 0.64<br>Reelin: p = 0.59<br>Sex: p = 0.01<br>CORT:Reelin: p = 0.41<br>CORT:Sex: p = 0.52<br>Reelin:Sex: p = 0.63<br>CORT:Reelin:Sex: p = 0.77 | F(1,48) = 0.23<br>F(2,48) = 0.53<br>F(1,48) = 6.79<br>F(1,48) = 0.69<br>F(1,48) = 0.43<br>F(2,48) = 0.47<br>F(1,48) = 0.09 | Marginal R^2 = 0.04<br>Conditional R^2 = 0.17 | Vehicle/Vehicle - Vehicle/Reelin: p = 0.07<br>Vehicle/Vehicle - CORT/Vehicle: p = 0.11<br>Vehicle/Vehicle - CORT/Reelin: p = 0.02<br>Vehicle/Vehicle - CORT/Repeated Reelin: p = 0.07<br>CORT/Vehicle - Vehicle/Reelin: p = 0.04<br>CORT/Vehicle - CORT/Reelin: p = 0.09<br>CORT/Vehicle - CORT/Repeated Reelin: p = 0.04<br>CORT/Reelin - CORT/Repeated Reelin: p = 0.05 | Vehicle/Vehicle - Vehicle/Reelin: p = 0.15<br>Vehicle/Vehicle - CORT/Vehicle: p < 0.001<br>Vehicle/Vehicle - CORT/Reelin: p = 0.20<br>Vehicle/Vehicle - CORT/Repeated Reelin: p = 0.27<br>CORT/Vehicle - Vehicle/Reelin: p = 0.15<br>CORT/Vehicle - CORT/Reelin: p = 0.20<br>CORT/Vehicle - CORT/Repeated Reelin: p = 0.27<br>CORT/Reelin - CORT/Repeated Reelin: p = 0.47 |
| 5De | Microglia process tracings - Total Nodes | Nested three-way ANOVA<br><br>Post hoc: Šídák's multiple comparisons test | Vehicle/Vehicle = 6<br>Vehicle/Reelin = 6<br>CORT/Vehicle = 6<br>CORT/Reelin = 6<br>CORT/Repeated | CORT: p = 0.06<br>Reelin: p = 0.94<br>Sex: p = 0.03<br>CORT:Reelin: p = 0.40<br>CORT:Sex: p = 0.36<br>Reelin:Sex: p = 0.36<br>CORT:Reelin:Sex: p = 0.56 | F(1,48) = 3.67<br>F(2,48) = 0.06<br>F(1,48) = 5.31<br>F(1,48) = 0.73<br>F(1,48) = 0.85<br>F(2,48) = 1.06<br>F(1,48) = 0.34 | Marginal R^2 = 0.07<br>Conditional R^2 = 0.27 | Vehicle/Vehicle - Vehicle/Reelin: p = 0.14<br>Vehicle/Vehicle - CORT/Vehicle: p = 0.08<br>Vehicle/Vehicle - CORT/Reelin: p = 0.28<br>Vehicle/Vehicle - CORT/Repeated Reelin: p = 0.30<br>CORT/Vehicle - Vehicle/Reelin: p = 0.06<br>CORT/Vehicle - CORT/Reelin: p = 0.36<br>CORT/Vehicle - CORT/Repeated Reelin: p = 0.38<br>CORT/Reelin - CORT/Repeated Reelin: p = 0.02 | Vehicle/Vehicle - Vehicle/Reelin: p = 0.27<br>Vehicle/Vehicle - CORT/Vehicle: p = 0.44<br>Vehicle/Vehicle - CORT/Reelin: p = 0.27<br>Vehicle/Vehicle - CORT/Repeated Reelin: p = 0.09<br>CORT/Vehicle - Vehicle/Reelin: p = 0.70<br>CORT/Vehicle - CORT/Reelin: p = 0.17<br>CORT/Vehicle - CORT/Repeated Reelin: p = 0.34<br>CORT/Reelin - CORT/Repeated Reelin: p = 0.17 |

| Figure Number | Figure Description | Statistical Test | n | p value | F values | Partial Eta Squared | R Squared | Post-Hocs | Post-Hocs Females | Post-Hocs Males | Cohen's d Fem | Cohen's d Mal | Westfall d Females | Westfall d Males |
|---|---|---|---|---|---|---|---|---|---|---|---|---|---|---|
| 6A | Microglia outline tracings - Area - Polymorphic layer | Nested linear mixed-effects model<br>Post hoc: Šídák's multiple comparisons test; Westfall d | Vehicle/Vehicle = 6<br>Vehicle/Reelin = 6<br>CORT/Vehicle = 6<br>CORT/Reelin = 6<br>CORT/Repeated Reelin = 6 | CORT: p < 0.001<br>Reelin: p < 0.001<br>Sex: p = 0.72<br>CORT:Reelin: p = 0.95<br>CORT:Sex: p = 0.07<br>Reelin:Sex: p = 0.67<br>CORT:Reelin:Sex: p = 0.24 | F(1,48) = 62.83<br>F(2,48) = 52.02<br>F(1,48) = 0.13<br>F(1,48) = 0.004<br>F(1,48) = 3.41<br>F(2,48) = 0.40<br>F(1,48) = 1.41 | | Marginal R^2 = 0.40<br>Conditional R^2 = 0.55 | | Vehicle/Vehicle - Vehicle/Reelin: p = 0.03<br>Vehicle/Vehicle - CORT/Vehicle: p = 0.006<br>Vehicle/Vehicle - CORT/Reelin: p = 1<br>Vehicle/Vehicle - CORT/Repeated Reelin: p = 0.30<br>CORT/Vehicle - Vehicle/Reelin: p < 0.001<br>CORT/Vehicle - CORT/Reelin: p < 0.001<br>CORT/Vehicle - CORT/Repeated Reelin: p < 0.001<br>CORT/Reelin - CORT/Repeated Reelin: p = 0.74 | Vehicle/Vehicle - Vehicle/Reelin: p < 0.001<br>Vehicle/Vehicle - CORT/Vehicle: p < 0.001<br>Vehicle/Vehicle - CORT/Reelin: p = 1<br>Vehicle/Vehicle - CORT/Repeated Reelin: p = 0.60<br>CORT/Vehicle - Vehicle/Reelin: p < 0.001<br>CORT/Vehicle - CORT/Reelin: p = 0.003<br>CORT/Vehicle - CORT/Repeated Reelin: p < 0.001<br>CORT/Reelin - CORT/Repeated Reelin: p = 0.32 | | | 0.938982995<br>1.093626061<br>0.187006624<br>0.626822507<br>2.032609056<br>1.280632685<br>1.720448568<br>0.439815883 | 1.550933762<br>1.293846472<br>0.122504578<br>0.497906038<br>2.844780235<br>1.171341895<br>1.79175251<br>0.620410616 |
| 6B | Microglia outline tracings - Perimeter - Polymorphic layer | Nested linear mixed-effects model<br>Post hoc: Šídák's multiple comparisons test; Westfall d | Vehicle/Vehicle = 6<br>Vehicle/Reelin = 6<br>CORT/Vehicle = 6<br>CORT/Reelin = 6<br>CORT/Repeated Reelin = 6 | CORT: p < 0.001<br>Reelin: p < 0.001<br>Sex: p = 0.70<br>CORT:Reelin: p < 0.001<br>CORT:Sex: p = 0.65<br>Reelin:Sex: p = 0.90<br>CORT:Reelin:Sex: p = 0.36 | F(1,48) = 50.99<br>F(2,48) = 46.43<br>F(1,48) = 0.16<br>F(1,48) = 12.55<br>F(1,48) = 0.21<br>F(2,48) = 0.11<br>F(1,48) = 0.84 | | Marginal R^2 = 0.36<br>Conditional R^2 = 0.51 | | Vehicle/Vehicle - Vehicle/Reelin: p = 0.68<br>Vehicle/Vehicle - CORT/Vehicle: p < 0.001<br>Vehicle/Vehicle - CORT/Reelin: p = 1<br>Vehicle/Vehicle - CORT/Repeated Reelin: p = 0.46<br>CORT/Vehicle - Vehicle/Reelin: p < 0.001<br>CORT/Vehicle - CORT/Reelin: p < 0.001<br>CORT/Vehicle - CORT/Repeated Reelin: p < 0.001<br>CORT/Reelin - CORT/Repeated Reelin: p = 0.18 | Vehicle/Vehicle - Vehicle/Reelin: p = 0.97<br>Vehicle/Vehicle - CORT/Vehicle: p < 0.001<br>Vehicle/Vehicle - CORT/Reelin: p = 1<br>Vehicle/Vehicle - CORT/Repeated Reelin: p = 0.71<br>CORT/Vehicle - Vehicle/Reelin: p < 0.001<br>CORT/Vehicle - CORT/Reelin: p < 0.001<br>CORT/Vehicle - CORT/Repeated Reelin: p < 0.001<br>CORT/Reelin - CORT/Repeated Reelin: p = 0.28 | | | 0.45571256<br>1.384002241<br>0.146933442<br>0.542611545<br>1.839714802<br>1.237068799<br>1.926613786<br>0.689544987 | 0.282332822<br>1.792059452<br>0.184536567<br>0.44163244<br>2.074392274<br>1.607522885<br>2.233691892<br>0.626169007 |
| 6C | Microglia outline tracings - Circularity - Polymorphic layer | Nested linear mixed-effects model<br>Post hoc: Šídák's multiple comparisons test; Westfall d | Vehicle/Vehicle = 6<br>Vehicle/Reelin = 6<br>CORT/Vehicle = 6<br>CORT/Reelin = 6<br>CORT/Repeated Reelin = 6 | CORT: p = 0.006<br>Reelin: p < 0.001<br>Sex: p = 0.46<br>CORT:Reelin: p < 0.001<br>CORT:Sex: p = 0.28<br>Reelin:Sex: p = 1<br>CORT:Reelin:Sex: p = 0.06 | F(1,48) = 8.06<br>F(2,48) = 11.18<br>F(1,48) = 0.56<br>F(1,48) = 30.35<br>F(1,48) = 1.18<br>F(2,48) = 0.0002<br>F(1,48) = 3.73 | | Marginal R^2 = 0.17<br>Conditional R^2 = 0.31 | | Vehicle/Vehicle - Vehicle/Reelin: p = 0.95<br>Vehicle/Vehicle - CORT/Vehicle: p = 0.004<br>Vehicle/Vehicle - CORT/Reelin: p = 0.89<br>Vehicle/Vehicle - CORT/Repeated Reelin: p = 1<br>CORT/Vehicle - Vehicle/Reelin: p = 0.07<br>CORT/Vehicle - CORT/Reelin: p = 0.10<br>CORT/Vehicle - CORT/Repeated Reelin: p = 0.002<br>CORT/Reelin - CORT/Repeated Reelin: p = 0.73 | Vehicle/Vehicle - Vehicle/Reelin: p = 0.04<br>Vehicle/Vehicle - CORT/Vehicle: p < 0.001<br>Vehicle/Vehicle - CORT/Reelin: p = 1<br>Vehicle/Vehicle - CORT/Repeated Reelin: p = 1<br>CORT/Vehicle - Vehicle/Reelin: p = 0.60<br>CORT/Vehicle - CORT/Reelin: p < 0.001<br>CORT/Vehicle - CORT/Repeated Reelin: p < 0.001<br>CORT/Reelin - CORT/Repeated Reelin: p = 1 | | | 0.264439891<br>0.982269774<br>0.310068842<br>0.07158905<br>0.717829883<br>0.672200931<br>1.053858823<br>0.381657892 | 0.775044137<br>1.203078657<br>0.030160137<br>0.099204239<br>0.42803452<br>1.17291852<br>1.302282896<br>0.129364376 |
| 6D | Microglia outline tracings - Max Feret - Polymorphic layer | Nested linear mixed-effects model<br>Post hoc: Šídák's multiple comparisons test; Westfall d | Vehicle/Vehicle = 6<br>Vehicle/Reelin = 6<br>CORT/Vehicle = 6<br>CORT/Reelin = 6<br>CORT/Repeated Reelin = 6 | CORT: p < 0.001<br>Reelin: p < 0.001<br>Sex: p = 0.64<br>CORT:Reelin: p < 0.001<br>CORT:Sex: p = 0.33<br>Reelin:Sex: p = 0.49<br>CORT:Reelin:Sex: p = 0.70 | F(1,48) = 34.97<br>F(2,48) = 58.76<br>F(1,48) = 0.22<br>F(1,48) = 15.05<br>F(1,48) = 0.97<br>F(2,48) = 0.72<br>F(1,48) = 0.14 | | Marginal R^2 = 0.27<br>Conditional R^2 = 0.36 | | Vehicle/Vehicle - Vehicle/Reelin: p = 0.43<br>Vehicle/Vehicle - CORT/Vehicle: p < 0.001<br>Vehicle/Vehicle - CORT/Reelin: p = 0.91<br>Vehicle/Vehicle - CORT/Repeated Reelin: p = 0.14<br>CORT/Vehicle - Vehicle/Reelin: p < 0.001<br>CORT/Vehicle - CORT/Reelin: p < 0.001<br>CORT/Vehicle - CORT/Repeated Reelin: p < 0.001<br>CORT/Reelin - CORT/Repeated Reelin: p = 0.85 | Vehicle/Vehicle - Vehicle/Reelin: p = 0.29<br>Vehicle/Vehicle - CORT/Vehicle: p < 0.001<br>Vehicle/Vehicle - CORT/Reelin: p = 0.99<br>Vehicle/Vehicle - CORT/Repeated Reelin: p = 0.05<br>CORT/Vehicle - Vehicle/Reelin: p < 0.001<br>CORT/Vehicle - CORT/Reelin: p < 0.001<br>CORT/Vehicle - CORT/Repeated Reelin: p < 0.001<br>CORT/Reelin - CORT/Repeated Reelin: p = 0.31 | | | 0.424647826<br>0.961309032<br>0.261429087<br>0.552234378<br>1.385956858<br>1.22273812<br>1.51354341<br>0.29080529 | 0.476598687<br>1.27257586<br>0.17583445<br>0.644031328<br>1.749174547<br>1.44841031<br>1.916607188<br>0.468196878 |
| 6E | Microglia outline tracings - Aspect Ratio - Polymorphic layer | Nested linear mixed-effects model<br>Post hoc: Šídák's multiple comparisons test; Westfall d | Vehicle/Vehicle = 6<br>Vehicle/Reelin = 6<br>CORT/Vehicle = 6<br>CORT/Reelin = 6<br>CORT/Repeated Reelin = 6 | CORT: p < 0.001<br>Reelin: p = 0.03<br>Sex: p = 0.89<br>CORT:Reelin: p = 0.02<br>CORT:Sex: p = 0.76<br>Reelin:Sex: p = 0.58<br>CORT:Reelin:Sex: p = 0.94 | F(1,48) = 17.99<br>F(2,48) = 3.67<br>F(1,48) = 0.02<br>F(1,48) = 6.27<br>F(1,48) = 0.09<br>F(2,48) = 0.55<br>F(1,48) = 0.005 | | Marginal R^2 = 0.03<br>Conditional R^2 = 0.06 | | Vehicle/Vehicle - Vehicle/Reelin: p = 0.95<br>Vehicle/Vehicle - CORT/Vehicle: p = 0.007<br>Vehicle/Vehicle - CORT/Reelin: p = 0.34<br>Vehicle/Vehicle - CORT/Repeated Reelin: p = 0.99<br>CORT/Vehicle - Vehicle/Reelin: p = 0.11<br>CORT/Vehicle - CORT/Reelin: p = 0.65<br>CORT/Vehicle - CORT/Repeated Reelin: p = 0.06<br>CORT/Reelin - CORT/Repeated Reelin: p = 0.88 | Vehicle/Vehicle - Vehicle/Reelin: p = 1<br>Vehicle/Vehicle - CORT/Vehicle: p = 0.02<br>Vehicle/Vehicle - CORT/Reelin: p = 0.95<br>Vehicle/Vehicle - CORT/Repeated Reelin: p = 0.92<br>CORT/Vehicle - Vehicle/Reelin: p = 0.04<br>CORT/Vehicle - CORT/Reelin: p = 0.25<br>CORT/Vehicle - CORT/Repeated Reelin: p = 0.30<br>CORT/Reelin - CORT/Repeated Reelin: p = 1 | | | 0.157112228<br>0.551461746<br>0.309517614<br>0.123946853<br>0.394349518<br>0.241944132<br>0.427514893<br>0.185570761 | 0.042268378<br>0.492898062<br>0.158743312<br>0.173407534<br>0.450629684<br>0.33415475<br>0.319490528<br>0.014664222 |
| 6F | Microglia outline tracings - Roundness - Polymorphic layer | Nested linear mixed-effects model<br>Post hoc: Šídák's multiple comparisons test; Westfall d | Vehicle/Vehicle = 6<br>Vehicle/Reelin = 6<br>CORT/Vehicle = 6<br>CORT/Reelin = 6<br>CORT/Repeated Reelin = 6 | CORT: p = 0.001<br>Reelin: p = 0.15<br>Sex: p = 0.74<br>CORT:Reelin: p = 0.05<br>CORT:Sex: p = 0.86<br>Reelin:Sex: p = 0.83<br>CORT:Reelin:Sex: p = 0.89 | F(1,48) = 11.98<br>F(2,48) = 1.96<br>F(1,48) = 0.11<br>F(1,48) = 4.00<br>F(1,48) = 0.03<br>F(2,48) = 0.19<br>F(1,48) = 0.02 | | Marginal R^2 = 0.02<br>Conditional R^2 = 0.05 | | Vehicle/Vehicle - Vehicle/Reelin: p = 0.98<br>Vehicle/Vehicle - CORT/Vehicle: p = 0.04<br>Vehicle/Vehicle - CORT/Reelin: p = 0.62<br>Vehicle/Vehicle - CORT/Repeated Reelin: p = 0.99<br>CORT/Vehicle - Vehicle/Reelin: p = 0.32<br>CORT/Vehicle - CORT/Reelin: p = 0.84<br>CORT/Vehicle - CORT/Repeated Reelin: p = 0.29<br>CORT/Reelin - CORT/Repeated Reelin: p = 0.99 | Vehicle/Vehicle - Vehicle/Reelin: p = 1<br>Vehicle/Vehicle - CORT/Vehicle: p = 0.10<br>Vehicle/Vehicle - CORT/Reelin: p = 0.92<br>Vehicle/Vehicle - CORT/Repeated Reelin: p = 0.94<br>CORT/Vehicle - Vehicle/Reelin: p = 0.24<br>CORT/Vehicle - CORT/Reelin: p = 0.73<br>CORT/Vehicle - CORT/Repeated Reelin: p = 0.68<br>CORT/Reelin - CORT/Repeated Reelin: p = 1 | | | 0.128163423<br>0.429447171<br>0.238788217<br>0.119507407<br>0.301283748<br>0.190658954<br>0.309939764<br>0.11928081 | 0.05918721<br>0.382178978<br>0.165642614<br>0.154584667<br>0.322991768<br>0.216536364<br>0.227594311<br>0.011057947 |
| 6G | Microglia outline tracings - Solidity - Polymorphic layer | Nested linear mixed-effects model<br>Post hoc: Šídák's multiple comparisons test; Westfall d | Vehicle/Vehicle = 6<br>Vehicle/Reelin = 6<br>CORT/Vehicle = 6<br>CORT/Reelin = 6<br>CORT/Repeated Reelin = 6 | CORT: p < 0.001<br>Reelin: p < 0.001<br>Sex: p = 0.25<br>CORT:Reelin: p = 0.003<br>CORT:Sex: p = 0.01<br>Reelin:Sex: p = 0.66<br>CORT:Reelin:Sex: p = 0.04 | F(1,48) = 29.08<br>F(2,48) = 13.21<br>F(1,48) = 1.37<br>F(1,48) = 9.55<br>F(1,48) = 6.46<br>F(2,48) = 0.42<br>F(1,48) = 4.48 | | Marginal R^2 = 0.23<br>Conditional R^2 = 0.37 | | Vehicle/Vehicle - Vehicle/Reelin: p = 0.12<br>Vehicle/Vehicle - CORT/Vehicle: p = 0.97<br>Vehicle/Vehicle - CORT/Reelin: p = 1<br>Vehicle/Vehicle - CORT/Repeated Reelin: p = 1<br>CORT/Vehicle - Vehicle/Reelin: p = 0.009<br>CORT/Vehicle - CORT/Reelin: p = 0.67<br>CORT/Vehicle - CORT/Repeated Reelin: p = 0.70<br>CORT/Reelin - CORT/Repeated Reelin: p = 1 | Vehicle/Vehicle - Vehicle/Reelin: p < 0.001<br>Vehicle/Vehicle - CORT/Vehicle: p = 0.79<br>Vehicle/Vehicle - CORT/Reelin: p = 0.94<br>Vehicle/Vehicle - CORT/Repeated Reelin: p = 1<br>CORT/Vehicle - Vehicle/Reelin: p < 0.001<br>CORT/Vehicle - CORT/Reelin: p = 1<br>CORT/Vehicle - CORT/Repeated Reelin: p = 0.89<br>CORT/Reelin - CORT/Repeated Reelin: p = 0.98 | | | 0.684047901<br>0.255315484<br>0.163667105<br>0.152617301<br>0.939363385<br>0.418982589<br>0.407932785<br>0.011049805 | 1.498353498<br>0.371273433<br>0.289114886<br>0.046279181<br>1.869626931<br>0.082158547<br>0.324994252<br>0.242835705 |
| 7A | Microglia outline tracings - Area - Subgranular zone | Nested linear mixed-effects model<br>Post hoc: Šídák's multiple comparisons test; Westfall d | Vehicle/Vehicle = 6<br>Vehicle/Reelin = 6<br>CORT/Vehicle = 6<br>CORT/Reelin = 6<br>CORT/Repeated Reelin = 6 | CORT: p = 0.27<br>Reelin: p = 0.40<br>Sex: p = 0.30<br>CORT:Reelin: p = 0.47<br>CORT:Sex: p = 0.31<br>Reelin:Sex: p = 0.63<br>CORT:Reelin:Sex: p = 0.03 | F(1,48) = 1.26<br>F(2,48) = 0.92<br>F(1,48) = 1.08<br>F(1,48) = 0.53<br>F(1,48) = 1.07<br>F(2,48) = 0.46<br>F(1,48) = 4.77 | | Marginal R^2 = 0.10<br>Conditional R^2 = 0.52 | | Vehicle/Vehicle - Vehicle/Reelin: p = 0.97<br>Vehicle/Vehicle - CORT/Vehicle: p = 0.46<br>Vehicle/Vehicle - CORT/Reelin: p = 0.87<br>Vehicle/Vehicle - CORT/Repeated Reelin: p = 0.60<br>CORT/Vehicle - Vehicle/Reelin: p = 0.98<br>CORT/Vehicle - CORT/Reelin: p = 1<br>CORT/Vehicle - CORT/Repeated Reelin: p = 1<br>CORT/Reelin - CORT/Repeated Reelin: p = 1 | Vehicle/Vehicle - Vehicle/Reelin: p = 1<br>Vehicle/Vehicle - CORT/Vehicle: p = 0.75<br>Vehicle/Vehicle - CORT/Reelin: p = 0.92<br>Vehicle/Vehicle - CORT/Repeated Reelin: p = 1<br>CORT/Vehicle - Vehicle/Reelin: p = 0.95<br>CORT/Vehicle - CORT/Reelin: p = 0.11<br>CORT/Vehicle - CORT/Repeated Reelin: p = 0.44<br>CORT/Reelin - CORT/Repeated Reelin: p = 1 | | | 0.373251289<br>0.737402204<br>0.516947049<br>0.660987701<br>0.364150915<br>0.220455154<br>0.076414503<br>0.144040652 | 0.166626352<br>0.577072175<br>0.445885636<br>0.16846928<br>0.410445823<br>1.022957811<br>0.745541455<br>0.277416356 |
| 7B | Microglia outline tracings - Perimeter - Subgranular zone | Nested linear mixed-effects model<br>Post hoc: Šídák's multiple comparisons test; Westfall d | Vehicle/Vehicle = 6<br>Vehicle/Reelin = 6<br>CORT/Vehicle = 6<br>CORT/Reelin = 6<br>CORT/Repeated Reelin = 6 | CORT: p = 0.20<br>Reelin: p = 0.20<br>Sex: p = 0.09<br>CORT:Reelin: p = 0.75<br>CORT:Sex: p = 0.24<br>Reelin:Sex: p = 0.75<br>CORT:Reelin:Sex: p = 0.67 | F(1,48) = 1.71<br>F(2,48) = 1.67<br>F(1,48) = 3.01<br>F(1,48) = 0.10<br>F(1,48) = 1.39<br>F(2,48) = 0.29<br>F(1,48) = 0.18 | | Marginal R^2 = 0.31<br>Conditional R^2 = 0.24 | | | | | | | |
| 7C | Microglia outline tracings - Circularity - Subgranular zone | Nested linear mixed-effects model<br>Post hoc: Šídák's multiple comparisons test; Westfall d | Vehicle/Vehicle = 6<br>Vehicle/Reelin = 6<br>CORT/Vehicle = 6<br>CORT/Reelin = 6<br>CORT/Repeated Reelin = 6 | CORT: p = 0.88<br>Reelin: p = 0.11<br>Sex: p = 0.52<br>CORT:Reelin: p = 0.80<br>CORT:Sex: p = 0.03<br>Reelin:Sex: p = 0.68<br>CORT:Reelin:Sex: p = 0.008 | F(1,48) = 0.02<br>F(2,48) = 2.32<br>F(1,48) = 0.41<br>F(1,48) = 0.07<br>F(1,48) = 4.93<br>F(2,48) = 0.38<br>F(1,48) = 7.70 | | Marginal R^2 = 0.09<br>Conditional R^2 = 0.37 | | Vehicle/Vehicle - Vehicle/Reelin: p = 0.53<br>Vehicle/Vehicle - CORT/Vehicle: p = 0.18<br>Vehicle/Vehicle - CORT/Reelin: p = 0.75<br>Vehicle/Vehicle - CORT/Repeated Reelin: p = 0.09<br>CORT/Vehicle - Vehicle/Reelin: p = 1<br>CORT/Vehicle - CORT/Reelin: p = 0.99<br>CORT/Vehicle - CORT/Repeated Reelin: p = 1<br>CORT/Reelin - CORT/Repeated Reelin: p = 0.93 | Vehicle/Vehicle - Vehicle/Reelin: p = 0.86<br>Vehicle/Vehicle - CORT/Vehicle: p = 0.07<br>Vehicle/Vehicle - CORT/Reelin: p = 0.97<br>Vehicle/Vehicle - CORT/Repeated Reelin: p = 1<br>CORT/Vehicle - Vehicle/Reelin: p = 0.71<br>CORT/Vehicle - CORT/Reelin: p = 0.46<br>CORT/Vehicle - CORT/Repeated Reelin: p = 0.08<br>CORT/Reelin - CORT/Repeated Reelin: p = 0.98 | | | 0.576255802<br>0.77560599<br>0.499566108<br>0.874221717<br>0.199350187<br>0.276039882<br>0.098615728<br>0.37465561 | 0.417381094<br>0.916059711<br>0.305875377<br>0.016631386<br>0.498678617<br>0.610184334<br>0.899428325<br>0.289243991 |
| 8 | Soma area | Nested linear mixed-effects model<br>Post hoc: Šídák's multiple comparisons test; Westfall d | Vehicle/Vehicle = 6<br>Vehicle/Reelin = 6<br>CORT/Vehicle = 6<br>CORT/Reelin = 6<br>CORT/Repeated Reelin = 6 | CORT: p = 0.29<br>Reelin: p = 0.87<br>Sex: p = 0.54<br>CORT:Reelin: p = 0.25<br>CORT:Sex: p = 0.13<br>Reelin:Sex: p = 0.33<br>CORT:Reelin:Sex: p = 0.15 | F(1,48) = 1.15<br>F(2,48) = 0.14<br>F(1,48) = 0.37<br>F(1,48) = 1.33<br>F(1,48) = 2.34<br>F(2,48) = 1.12<br>F(1,48) = 2.10 | | Marginal R^2 = 0.03<br>Conditional R^2 = 0.21 | | | | | | | |

| Figure Number | Figure Description | Statistical Test | n | p value | F values | Partial Eta Squared | R Squared | Post-Hocs | Post-Hocs Females | Post-Hocs Males | Cohen's d Fem | Cohen's d Mal | Westfall d Females | Westfall d Males |
|---|---|---|---|---|---|---|---|---|---|---|---|---|---|---|
| 9A | Cleaved caspase-3 immunoreactivity - Subgranular zone | Three-way ANOVA | Vehicle/Vehicle = 6 | CORT: p = 0.03 | F(1,48) = 4.98 | 0.090540336 | | | Vehicle/Vehicle - Vehicle/Reelin: p = 0.97 | Vehicle/Vehicle - Vehicle/Reelin: p = 1 | 0.8 | 0.02 | | |
| | | Post hoc: | Vehicle/Reelin = 6 | Reelin: p = 0.01 | F(2,48) = 4.99 | 0.166519708 | | | Vehicle/Vehicle - CORT/Vehicle: p = 0.01 | Vehicle/Vehicle - CORT/Vehicle: p = 0.94 | 1.3 | 0.32 | | |
| | | Šidák's multiple comparisons test | CORT/Vehicle = 6 | Sex: p = 0.20 | F(1,48) = 1.66 | 0.032119207 | | | Vehicle/Vehicle - CORT/Reelin: p = 1 | Vehicle/Vehicle - CORT/Reelin: p = 0.98 | 0.93 | 0.7 | | |
| | | | CORT/Reelin = 6 | CORT:Reelin: p = 0.16 | F(1,48) = 1.99 | 0.038298326 | | | Vehicle/Vehicle - CORT/Repeated Reelin: p = 1 | Vehicle/Vehicle - CORT/Repeated Reelin: p = 1 | 1.03 | 0.1 | | |
| | | | CORT/Repeated Reelin = 6 | CORT:Sex: p = 0.24 | F(1,48) = 1.44 | 0.027957558 | | | CORT/Vehicle - Vehicle/Reelin: p < 0.001 | CORT/Vehicle - Vehicle/Reelin: p = 1 | 2.1 | 0.34 | | |
| | | | | Reelin:Sex: p = 0.05 | F(2,48) = 3.12 | 0.11108507 | | | CORT/Vehicle - CORT/Reelin: p = 0.03 | CORT/Vehicle - CORT/Reelin: p = 1 | 2.23 | 0.38 | | |
| | | | | CORT:Reelin:Sex: p = 0.47 | F(1,48) = 0.53 | 0.010391945 | | | CORT/Vehicle - CORT/Repeated Reelin: p = 0.002 | CORT/Vehicle - CORT/Repeated Reelin: p = 0.98 | 2.33 | 0.42 | | |
| | | | | | | | | | CORT/Reelin - CORT/Repeated Reelin: p = 0.99 | CORT/Reelin - CORT/Repeated Reelin: p = 1 | 0.1 | 0.8 | | |
| 9B | Cleaved caspase-3 immunoreactivity - Polymorphic layer | Three-way ANOVA | Vehicle/Vehicle = 6 | CORT: p = 0.36 | F(1,48) = 0.87 | 0.017061448 | | | Vehicle/Vehicle - Vehicle/Reelin: p = 0.78 | Vehicle/Vehicle - Vehicle/Reelin: p = 1 | 0.54 | 0.28 | | |
| | | Post hoc: | Vehicle/Reelin = 6 | Reelin: p = 0.005 | F(2,48) = 6.02 | 0.194118716 | | | Vehicle/Vehicle - CORT/Vehicle: p = 0.21 | Vehicle/Vehicle - CORT/Vehicle: p = 1 | 1.97 | 0.62 | | |
| | | Šidák's multiple comparisons test | CORT/Vehicle = 6 | Sex: p = 0.52 | F(1,48) = 0.42 | 0.008269893 | | | Vehicle/Vehicle - CORT/Reelin: p = 0.61 | Vehicle/Vehicle - CORT/Reelin: p = 0.88 | 0.2 | 0.5 | | |
| | | | CORT/Reelin = 6 | CORT:Reelin: p = 0.37 | F(1,48) = 0.82 | 0.016069017 | | | Vehicle/Vehicle - CORT/Repeated Reelin: p = 0.48 | Vehicle/Vehicle - CORT/Repeated Reelin: p = 1 | 0.27 | 0.11 | | |
| | | | CORT/Repeated Reelin = 6 | CORT:Sex: p = 0.79 | F(1,48) = 0.07 | 0.00138698 | | | CORT/Vehicle - Vehicle/Reelin: p = 0.005 | CORT/Vehicle - Vehicle/Reelin: p = 1 | 2.51 | 0.34 | | |
| | | | | Reelin:Sex: p = 0.01 | F(2,48) = 4.67 | 0.157484571 | | | CORT/Vehicle - CORT/Reelin: p = 0.003 | CORT/Vehicle - CORT/Reelin: p = 1 | 1.77 | 0.12 | | |
| | | | | CORT:Reelin:Sex: p = 0.12 | F(1,48) = 2.51 | 0.047801481 | | | CORT/Vehicle - CORT/Repeated Reelin: p = 0.001 | CORT/Vehicle - CORT/Repeated Reelin: p = 0.99 | 2.25 | 0.5 | | |
| | | | | | | | | | CORT/Reelin - CORT/Repeated Reelin: p = 1 | CORT/Reelin - CORT/Repeated Reelin: p = 0.78 | 0.47 | 0.38 | | |
